# Endothelial miR-34a deletion guards against aneurysm development despite endothelial dysfunction

**DOI:** 10.1101/2024.09.12.612788

**Authors:** Aleksandra Kopacz, Damian Kloska, Anna Bar, Marta Targosz-Korecka, Dominik Cysewski, Stefan Chlopicki, Alicja Jozkowicz, Anna Grochot-Przeczek

## Abstract

**Objectives:** Our previous study reported a reciprocal link between NRF2, a stress-responsive cytoprotective transcription factor, and aortic and endothelial cell (EC) ageing. We also found that NRF2 transcriptional knockout (tKO) mice are prone to abdominal aortic aneurysm (AAA) development. Given that miRNA-34a is a marker of ageing, in this study we explored its relationship with NRF2 and its role in vascular function and AAA formation.

**Approach and results:** The experiments were performed in primary human aortic endothelial cells (HAECs) from young and aged donors and mice devoid of NRF2 transcriptional activity and endothelial miR-34a. The normolipidemic mice were challenged with angiotensin II (Ang II) to develop AAA. We show that premature NRF2-dependent aging of aortic endothelial cells (ECs) depends on miR-34a. Infusion of hypertensive Ang II in mice increases miR-34a in the aortic endothelial layer and serum, especially in mice which develop AAA. Mice deficient in endothelial miR-34a (miR-34a^ΔEC^) display severe EC dysfunction. Despite that, such mice are protected from AAA development, also on the NRF2 tKO background. Ang II infusion increases proliferation of intimal ECs in these mice. The protective effect of endothelial miR-34a deletion on AAA formation is reversed by rapamycin that suppresses EC proliferation. MTA2, but not SIRT1, is a direct target of miR-34a abrogating Ang II-induced EC proliferation.

**Conclusions:** These findings reveal that AAA development in NRF2 tKO mice relies on endothelial miR-34a overexpression. Deletion of endothelial miR-34a protects mice from AAA despite inducing endothelial cell dysfunction. The fine-tuning of EC proliferation may play a therapeutic role in the treatment of aneurysm.

## Introduction

An abdominal aortic aneurysm (AAA), local, usually infrarenal dilation of the aorta, is a complex and fatal vascular disease, for which surgery remains the only therapeutic option.^1,2^ Several risk factors for the development of AAA such as male sex, ageing, smoking, hypertension, and family history, are outlined. Among general characteristics of AAA are increased degradation of extracellular matrix, immune cell recruitment, loss of vascular smooth muscle cells (VSMC), as well as media and adventitia thinning.^1,2^ However, detailed activation pathways, which alter the architecture of the aortic wall, are still not clear.

Furthermore, the significance of endothelial cells (ECs), which line all blood vessels and form the intimal layer of the aorta, in the development of AAA is generally unknown. Notably, endothelium governs the proper functioning of the cardiovascular system, producing vasodilatory nitric oxide (NO), and maintaining the anti-thrombotic and anti-inflammatory phenotype of blood vessels.^3^ ECs are very vulnerable, and exposing them to adverse signals can trigger their premature ageing and dysfunction. Subsequently, this impairs blood vessel function and increases the risk of cardiovascular diseases.^4^

Recently, we showed that mice lacking NRF2 (nuclear factor (erythroid-derived 2)-like 2) transcriptional activity are more prone to AAA development compared to wild type (WT) counterparts.^5^ NRF2 is a stress-responsive protein, the major coordinator of cellular response to oxidative stress, which transactivates the expression of numerous detoxifying, antioxidative and protective genes.^6,7^ NRF2 plays a critical role in ageing.^4,8^ Although the extent of the deterioration of NRF2 signalling differs between tissues, for the cardiovascular system, there is a concise age-dependent decrease, as shown in rat aortas,^9^ monkey carotid arteries,^10^ and also human aortic endothelial cells (HAECs).^11^ Furthermore, NRF2 knockdown triggers premature senescence of human ECs *in vitro*^12^ and the aortas of mice lacking transcriptional activity of NRF2 undergo premature ageing.^13^ Given the important role of NRF2 in ECs and strong reciprocal associations of NRF2, ageing and cardiovascular diseases, we hypothesised that the molecular determinant of the protective effect of NRF2 in AAA could be the endothelial age-related factor.

A hallmark molecule overexpressed in senescent cells, including epithelial, endothelial, vascular smooth muscle, and cancer cells, is miRNA-34a (miR-34a).^14–17^ Its levels are also elevated in aged hearts and aortas.^14,18^ It is worth to note, that although genetic ablation of miR-34a does not cause obvious developmental or pathological abnormalities, and the knockout mice are born at the expected Mendelian ratio,^19^ the aortas of miR-34a KO mice are characterised by increased medial thickness and cellular density.^14^

Importantly, miR-34a expression is higher within the aneurysm compared to healthy human aorta,^20^ and in peripheral blood mononuclear cells (PBMCs)^21^ of patients with AAA compared to control donors. This is not a correlation only but also a causative relation, as AAV9-mediated inhibition of miR-34a in hyperlipidaemic ApoE KO mice revealed a protective effect of miR-34a decrease on angiotensin II (Ang II)-induced AAA formation.^20^ The authors of this report postulate the regulation of VSMC senescence dependent on METTL3/miR-34a/SIRT1 as a molecular mechanism.^20^ In contrast, our preliminary data indicate that strong upregulation of miR-34a induced by Ang II is specific for endothelial cells. Therefore, in this study, we took advantage of the conditional endothelial miR-34a knockout mouse model to address the question of the role of endothelial miR-34a in the AAA formation, in the context of ageing and NRF2-dependent premature senescence.

## Materials and methods

### Animals

Mice were kept in a constant 14/10 h light/dark cycle at an environmental temperature of 22 ± 2°C and provided with a food and water *ad libitum*, except for special procedures as indicated in the detailed description of the method. For all animal studies, mice were handled strictly in accordance with the good animal practices, following the guidelines of Directive 2010/63/EU of the European Parliament on the protection of animals used for scientific purposes, local ethics committee approvals, and the institutional recommendations. We used EC-specific knockout of miR-34a, which was obtained by crossing miR-34a floxed mice (Mir34atm1.2Aven/J, background: C57BL/6 x 129S4/SvJae, strain: #018545; Jackson Laboratories) with Cdh5 (PAC)-CreERT2 mice (strain #13073, background: C57BL/6, Cancer Research Technology Repository at Taconic; developed by Dr Ralf Adams) and NRF2 transcriptional knockout (tKO) mice (background: C57BL/6), initially developed by Prof. Masayuki Yamamoto (Tohoku University, Japan), kindly provided by Prof. Antonio Cuadrado (Universidad Autonoma de Madrid, Spain). In these mice, the NRF2 DNA binding domain is replaced by LacZ gene,^12,22^ which abrogates NRF2 transcriptional activity. The presence of the NRF2-LacZ fusion protein does not have an effect on the detection of senescence-associated β-galactosidase (SA-β-gal).^13^

Three-month and six-month-old mice were used in the study. For the experiments with Ang II infusion only males were taken. All other experiments were conducted on male/female mice in a 1:1 ratio. No sex-related differences were found in the parameters analysed. The procedures were accepted by the II Local Ethics Committee in Kraków 283/2020, 320/2020, 16/2021, 166/2021, 171/2021, 249/2021, 349/2021, 273/2022 and annexes to these approvals. For breeding, the donor of Cre recombinase was always a male and, if mice were crossed further, it was possible up to fifth generation (F5). For final comparisons, we always compared Cre recombinase-bearing mice, which differed at the floxed gene site, to minimise the potential bias related to the effect of tamoxifen on aortic function.^23^ In order to induce Cre-loxP recombination, mice were injected intraperitoneally (*i.p.*) with tamoxifen (75 mg/kg in corn oil, Sigma-Aldrich) for the consecutive 5 days. Mice were analysed 10 days after the last dose of tamoxifen, except for aneurysm studies, where the withdrawal period lasted 30 days. For induction of AAA, mice were infused via osmotic minipumps (Alzet) with Ang II (1000 ng/min/kg, Bachem) or saline for up to 14 days. Rapamycin (4 mg/kg, Selleckchem) together with 5-ethynyl-2’-deoxyuridine (EdU, 50 mg/kg, ThermoFisher) diluted in 5% Tween 80, 5% Kolliphor, were injected *i.p.* daily, for 7 days. All procedures requiring anaesthesia, described in details in the supplementary methods, were carried out using ketamine (100 mg/kg) and xylazine (10 mg/kg) administered *i.p.* once or isoflurane 1.5 vol% in oxygen and air (2:3) mixture, inhaled for maximum 30 min. Anaesthesia was monitored by the loss of pedal reflex. Mice were euthanized by cervical dislocation or cutting the spinal cord for procedures requiring anaesthesia or using carbon dioxide if the animals were sacrificed for tissue collection.

### Cell culture

Human aortic endothelial cells (HAECs) isolated from young and aged donors under informed consent were purchased from Gibco, Life Technologies and Promocell. The investigations conform to the principles outlined in the Declaration of Helsinki. HAECs were grown in Endothelial Basal Medium (EBM-2) (Lonza) supplemented with EGM-2MV SingleQuot Kit Supplements & Growth Factors without antibiotic (Lonza) and with 5% foetal bovine serum (FBS, EURx). Cells were grown at 37°C under a humidified incubator in 5% CO_2_ atmosphere. The cells used in all experiments were between passages 5 and 14 to exclude the impact of senescence caused by prolonged cell culture, which was monitored by SA-βgal activity. The cells were stimulated with 500 nM Ang II for 6 h in EGM-2MV medium.

### Blood pressure measurement

Mice underwent non-invasive blood pressure measurement by tail-cuff plethysmography (BP-2000 series II, Visitech Systems) after a period of adaptation and before the commencement of the treatment protocol. The results were presented as a percentage change compared to day 0 (after adaptation time and before the administration of any drugs).

### Intravital assessment of leukocyte adhesion

4 h prior to the experiment, mice were injected with tumor necrosis factor-α (TNFα, 2 µg/kg, R&D). After that time, mice were anaesthetized with ketamine and xylazine (100 mg/kg and 10 mg/kg, respectively). Upon confirmation of loss of pedal reflex, mice were injected intravenously with rhodamine 6G (1 mg/kg, Sigma-Aldrich). After 1 minute, the abdominal cavity was opened and the mesenteric venules visualized gently, using cotton buds. The mouse was placed in the fluorescent microscope (Nikon) and the flow of leukocytes was recorded for 1 minute in at least 3 parts of the mesentery. After the procedure, mice were euthanized by cervical dislocation.

### Induction of thrombosis by extrinsic pathways (collagen – epinephrine)

4 h before the experiment, mice were injected with TNFα (2 µg/kg, R&D). After that time, mice were anaesthetized with ketamine and xylazine (100 mg/kg and 10 mg/kg, respectively). Upon confirmation of the loss of pedal reflex, mice were injected with 10 µg of Alexa Fluor (AF) 488-labeled anti-CD41 antibody (Biolegend) into the retroorbital sinus. After 5 minutes, mice were injected with collagen and epinephrine to induce thrombosis. After exactly 5 minutes, mice were euthanized by cutting the spinal cord and the vasculature was perfused through the right ventricle with 10 mL of 4% paraformaldehyde (PFA). Lungs were collected for subsequent analyses.

### Mass spectrometry

Proteome profiling of HAECs was performed according to the protocol published in ^12^.

### Pulse wave velocity (PWV)

Measurements were performed using a Doppler flow velocity system (Indus Instruments, Scintica Instrumentation) as described previously.^24^ During Doppler measurements, mice were anaesthetized using isoflurane (Aerrane; Baxter, 1.5 vol% in oxygen and air (2:3) mixture). PWV was measured using two 20-MHz Doppler probes, simultaneously recording velocity signals from 2 sites, representative of the thoracic and abdominal aorta. The aortic PWV was calculated by dividing the separation distance by the difference in arrival times of the velocity pulse timed with respect to the ECG. Signal analysis was performed using a Doppler signal processing workstation (version 1.625, Indus Instruments).

### Atomic force microscopy (AFM) nanoindentation

After removal of perivascular fat, the abdominal aorta was cut into small patches to expose the inner wall of the vessel. The aortas were prepared as for *en face* staining and glued to the glass using CellTak (BD Biosciences). After preparation, the samples were placed in Krebs-HEPES buffer pH 7.4 (100 mM NaCl, 4.7 mM KCl, 1.2 mM MgSO_4_, 1 mM KH_2_PO_4_, 2.5 mM CaCl_2_, 25 mM NaHCO_3_, 5.6 mM glucose, 20 mM HEPES) for 1 h. The buffer was changed to Hanks’ balanced salt solution (HBSS) supplemented with 1% foetal bovine serum (FBS, EURx), 100 U/mL penicillin, 100 µg/mL streptomycin, and 5 mM glucose. If not specified, all of the reagents were from Sigma-Aldrich. AFM nanoindentation experiments were performed with a NanoWizard III system (JPK, Germany). We used a spherical colloidal probe (4.5 μm diameter, Novascan), attached to a cantilever with a spring constant of 0.03 N/m. Indentation curves were recorded for a maximum loading force of 1 nN at a velocity of 1.5 μm/s. For each aorta sample, spatial maps of indentation curves were recorded at many random positions of the sample. Typically, each region of interest comprised 100 curves that were recorded in a 20×20 μm area. If necessary, for better resolution, the size of the grids was increased. The analysis was performed following the protocol published elsewhere.^25^ For the analysis of glycocalyx, Krebs-HEPES buffer was supplemented with 2% serum collected from the analysed mouse. The extraction of different parameters (corresponding to different parts and layers of the aorta) was based on the range of nanoindentation.

### *En face* immunofluorescent staining

For *en face* immunostaining, mice were anesthetised with ketamine and xylazine (100 mg/kg and 10 mg/kg, respectively) and perfused with 4% PFA. The entire aorta was removed, gently cleaned of perivascular fat, and fixed for 15 minutes with 4% PFA. The aortas were incubated for 3 h in blocking buffer (5% goat serum, 1% BSA in PBS) and then overnight with the antibodies (AF647-conjugated anti-CD31, M.EC. 13.3, BioLegend; ATTO-550-conjugated anti-AT1R antibody, Allomone labs) or Wheat Germ Agglutinin (WGA), Rhodamine conjugate (Vector Labs) diluted in 3% BSA. After that, the aortas were washed 3 times, and, if needed, incubated with the host-specific antibodies conjugated with Alexa Fluor fluorophores (dilution 1:1000; IgG H+L, Life Technologies) and DAPI (1 µg/mL, Sigma-Aldrich) for 2 h at room temperature. After 3 series of washing, the aortas were opened, the intima facing up, and mounted with Dako Fluorescence Mounting Medium (Dako). To visualise EdU positive cells Click-iT (Invitrogen) was used. High-resolution images were taken using a meta laser scanning confocal microscope (LSM-880; Carl Zeiss). By modulating the Z-axis, we could focus on different types of cells. The final distinction was made on the basis of the nuclear shape, round for endothelial cells, and significantly elongated for smooth muscle cells.

### Detection of SA-β-gal activity in aortas

The aortas were fixed for 3 minutes with 4% formalin, washed twice with PBS and incubated overnight with a staining solution (5 mM potassium ferricyanide, 5 mM potassium ferrocyanide, 150 mM NaCl, 2 mM MgCl_2_, 1 mg/mL X-gal in citrate buffer pH 6). The next day, the aortas were washed with PBS. For the depiction of the vascular layer, the aortas were embedded in OCT freezing medium, frozen, and the aortic specimens were cut on cryostat for further immunofluorescent staining with CD31, αSMA (Abcam) antibodies. Images were taken in the brightfield and fluorescence using a confocal microscope LSM 880 (Zeiss).

### *In situ* hybridization

Aortas (or aortic specimens cut on cryostat to 10 µm) were fixed in 4% PFA for 10 minutes, washed with PBS, digested for 8 minutes with 20 µg/mL proteinase K in digestion buffer (50 mM Tris HCl pH 7.4, 1 mM EDTA pH 8.0), followed by 5 minutes incubation in 0.2% glycine in PBS (Bio-shop). After PBS wash, the aortas were refixed for 10 minutes with 4% PFA, and then washed twice with PBS. For the acetylation step, the aortas were treated for 10 minutes first with 0.1 M triethanolamine pH 8 (Sigma-Aldrich), then with 0.1 M triethanolamine with 0.25% acetic acid, which was followed by PBS wash twice. For the hybridisation step, the stock 3’DIG-labelled probes (scrambled (included in the miRCURY LNA Control Set - Qiagen); miRNA-34a custom probe YD00618250-BEG) were diluted to 25 µM in hybridisation buffer (50% formamide, 5x SSC, 5x Denhardt’s solution, 500 mg/mL salmon sperm DNA, 250 mg/mL tRNA – all from ThermoFisher), denatured for 5 minutes in 85°C, and then, after cooling, placed on the aortas for the overnight hybridization at 50°C. The next day, the aortas were washed for 5 minutes at room temperature with 5x SSC, followed by a 30 minute wash with 20% formamide/0.5x SSC at hybridization temperature. Subsequently, the aortas were washed for 5 minutes with TBST buffer (0.1% Tween in TBS), then blocked with blocking buffer (10% sheep serum in TBST) for 1 h. The next steps of the protocol differed depending on the used aortic samples. Whole mount aortas were used for *en face* fluorescent staining, and incubated overnight with rhodamine-conjugated anti-DIG antibody (Roche, 1:500) and Alexa Fluor 647-coupled anti-CD31 antibody (clone MEC13.3, BioLegend, 1:100) diluted in blocking buffer. On the following day, the aortas were counterstained with DAPI, washed with PBS, opened longitudinally and mounted with Dako Fluorescence Mounting Medium (Dako). For conventional miRNA ISH protocols, the aortic sections were incubated overnight at 4°C in a water-humidified chamber with anti-DIG-AP antibody diluted in blocking buffer (Roche, 1:500), followed by a three-time wash with TBST on the next day. Then the slides were incubated in a staining solution at 32°C for 4 h, followed by wash in TBST and final fixation in 4% PFA for 10 minutes.

### miRNA isolation

miRNA was isolated from 400 µL of mouse serum using the miRNeasy Serum/Plasma Kit (Qiagen) according to the attached protocol. Total RNA from HAECs was isolated by phenol-chloroform extraction and precipitation in 50% isopropanol. The reverse transcription reaction was carried out using Mistiq microRNA and cDNA Synthesis Mix (Sigma-Aldrich) and following the vendor’s instructions. Quantitative RT-PCR was performed in a StepOnePlus real-time PCR system (Applied Biosystems) using SYBR Green PCR Master Mix (Sigma). Primer sequences for miR-34a-5p: 5’-TGGCAGTGTCTTAGCTGGTTGT −3’; U6 snRNA: 5’-CGCAAGGATGACACGCAAATC-3’.

### Assessment of endothelial NO production in the isolated aorta using electron paramagnetic resonance (EPR)

For measurements of NO, EPR spin-trapping with diethyldithiocarbamic acid sodium salt (DETC) was used. Briefly, Krebs–HEPES buffer (consisting of, in mM: NaCl 99.0; KCl 4.7; MgSO_4_·7H_2_O 1.2; KH_2_PO_4_ 1.0; CaCl_2_·H_2_O 2.5; NaHCO_3_ 25.0; glucose 5.6; and Na-HEPES 20.0) was filtered through a 0.22 µm paper syringe filter and equilibrated to pH 7.4, then deoxygenized by bubbling argon gas for at least 30 minutes. DETC (3.6 mg) and FeSO_4_ · 7H_2_O (2.25 mg) were separately dissolved under argon gas bubbling in two 10 mL volumes of ice-cold Krebs–HEPES buffer and were kept under gas flow on ice until used. Half (upper segment) of freshly harvested aortas was cleaned from adherent tissue, opened longitudinally, and placed in wells of the 48-well plate, each filled with 100 μL Krebs–HEPES and preincubated for 30 minutes at 37°C. The DETC and FeSO_4_ · 7H_2_O solutions were mixed 1:1 (v/v) to obtain 250 μL Fe(DETC)_2_ colloid per well (final concentration 285 μM) and immediately added to the aorta in parallel with calcium ionophore A23187 (final concentration 1 μM) to stimulate endothelial nitric oxide synthase (eNOS) and subsequently incubated at 37°C for 90 minutes. After incubation, each aorta was removed from the buffer, drained on a piece of kimwipe for 5 s, and the wet mass was measured. Next, the aorta was frozen in liquid nitrogen, into the middle of a 400 μL column of Krebs–HEPES buffer and stored in −80°C until measured.

### Transfection

HAECs were transfected either with 20 nM siRNA against human *NFE2L2* (Life Technologies, s9493), *TP53* (Life Technologies, s605), *SIRT1* (Life Technologies, s223591), *MTA2* (Thermo Scientific, 4392420) or scrambled siRNA (Life Technologies, 4390846) or 10 nM anti-miR-34a-5p (AM11030), pre-miR-34a-5p (AM17100) or neg anti-miR (AM17010) using Lipofectamine™ RNAiMAX Transfection Reagent (Life Technologies) in Opti-MEM I Reduced Serum medium (Life Technologies). The experiments were performed 48 h after transfection. The transfections of more than one target were performed concomitantly.

### Detection of SA-β-gal activity in HAECs

Cells were fixed for 3 minutes with 4% formaldehyde, washed twice with PBS, and incubated overnight with a staining solution (5 mM potassium ferricyanide, 5 mM potassium ferrocyanide, 150 mM NaCl, 2 mM MgCl_2_, 1 mg/mL X-gal in citrate buffer pH 6). The next day, cells were washed with PBS and stained with Hoechst 33342 to detect nuclei. Images were taken in the brightfield and fluorescence using a fluorescent microscope (Nikon).

### PCNA, p53 and MTA2 immunofluorescent staining

Cells were fixed for 8 minutes in 4% PFA, permeabilized with 0.2% Triton X-100 in PBS for 20 minutes, and blocked with 10% goat serum for 1 h. The cells were incubated with anti-PCNA (Life Technologies, MA511358), anti-p53 (Abcam, ab131442) or anti-MTA2 (Abcam, ab171073) antibody in 3% bovine serum albumin overnight at 4°C. The next day, the cells were incubated with a secondary Alexa Fluor 568-labelled anti-mouse antibody (Invitrogen) and DAPI to stain the nuclei.

### Angiotensin (1-7), IL-1β, IL-6 and sICAM measurement

Angiotensin (1-7) was measured by the Abbexa ELISA test, according to the vendor’s protocol. The levels of interleukins (IL-1β and IL-6) and soluble intercellular adhesion molecule 1 (sICAM) were assessed by ELISA tests from R&D.

### Measurement of APTT, PT, and fibrinogen

The coagulation parameters were checked by using Bio-ksel tests, according to the vendor’s protocol.

### Statistical analysis

All experiments were performed in duplicates and were repeated at least three times. Data are presented as mean ± SEM. Gaussian distribution of data was tested with the Shapiro-Wilk test. Statistical assessment was done with Mann-Whitney U test for two group comparisons or by analysis of variance (ANOVA), followed by a Tukey posthoc test for multiple comparisons. Grubbs’ test was used to detect outliers in data sets. Fisher’s exact test was used to calculate the frequency of aneurysm appearance and rupture. Differences were accepted as statistically significant for p < 0.05.

## Results

### The relevance of miR-34a in the premature and physiological ageing of human aortic endothelial cells

Previous studies have shown that miR-34a levels are elevated in aged mouse aorta^14^, aged human or mouse hearts^18^, and in human umbilical vein endothelial cells (HUVECs) subjected to replicative senescence^26^, whereas transfection with pre-miR-34a induced premature senescence of HUVEC.^16^ Similarly, both replicative and physiological ageing led to elevation of miR-34a in human aortic smooth muscle cells (HASMCs). However, no such data are available for human aortic endothelial cells, which govern healthy phenotype and function of the aorta.

To address this point, we examined miR-34a-5p (henceforth called miR-34a) expression in human aortic endothelial cell (HAEC) lines, three from young (21-23 years) and four from aged (62-89 years) donors. The level of miR-34a, normalised to the corresponding U6 snRNA, was significantly higher in aged donor-derived HAECs (Fig. 1A). It was accompanied by the increased level of p53 (Fig. 1B, C), the major factor transactivating miR-34a expression. On the other hand, upregulation of miR-34a in young HAECs transfected with pre-miR-34a (Fig. 1D) potently increased both the activity of senescence-associated β-galactosidase (SA-β-gal) (Fig. 1E) and the production of senescence-associated secretory phenotype (SASP) factors: IL-1β, IL-6 and sICAM-1 (Fig. 1F).

**Fig. 1.**
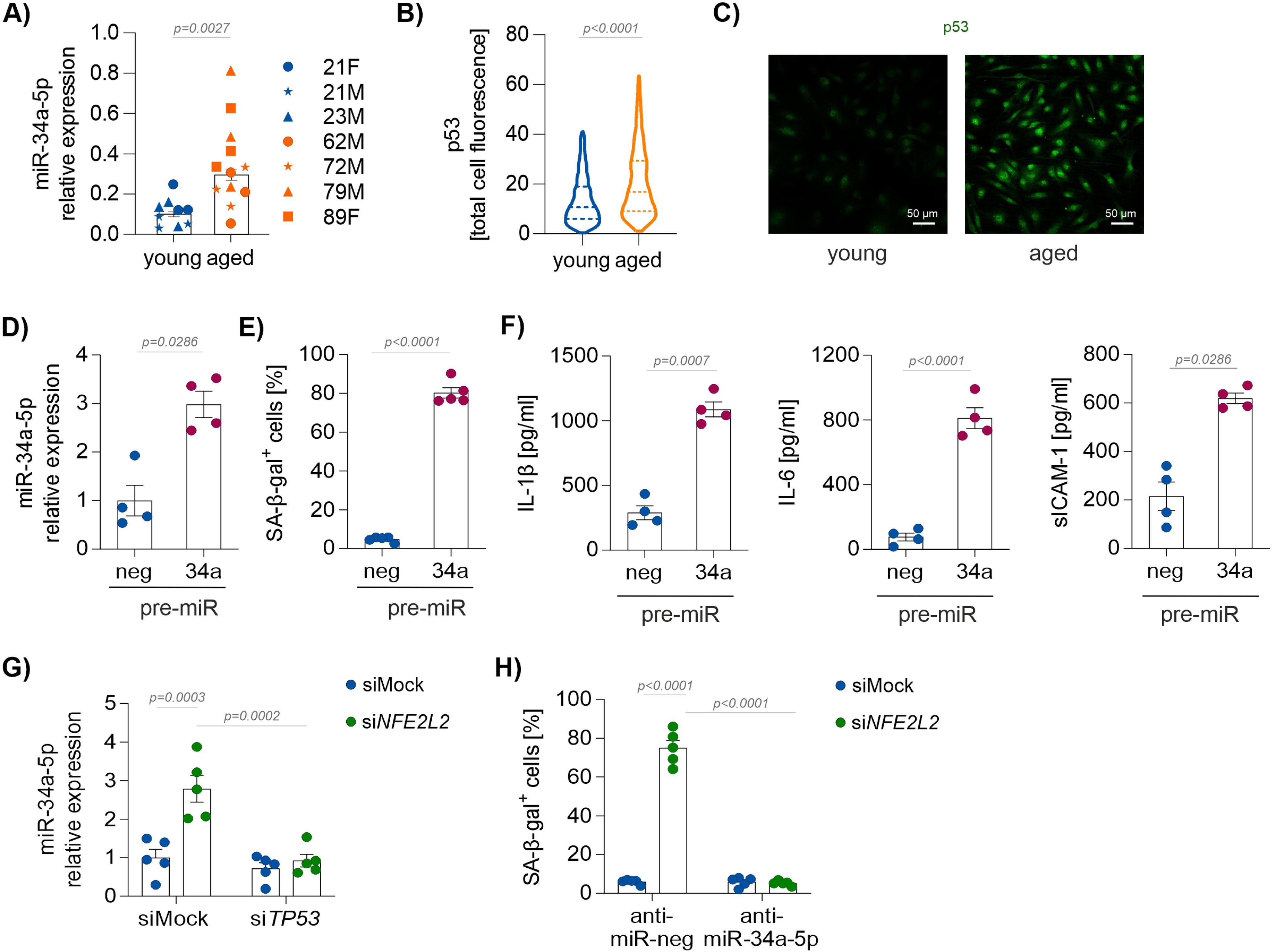
The crosstalk of miR-34a and NRF2 in ageing. **(A)** Expression of miR-34a-5p in HAECs derived from young and aged donors. Relative expression was measured by real-time PCR. U6 snRNA served as an internal control. n=9–12, Mann-Whitney U test. **(B,C)** Protein level of p53 in HAECs derived from 3 young and 5 aged donors. (B) Quantitative data and (C) representative pictures. Mann-Whitney U test. Scale bar 50 µm. **(D)** Expression of miR-34a-5p in young donor-derived HAECs transfected with pre-miR-34a or negative control. Relative expression was measured by real-time PCR. U6 snRNA served as an internal control. n=4, Mann-Whitney U test. **(E)** Percentage of cells with increased SA-β-gal activity in young donor-derived HAECs transfected with pre-miR-34a or negative control. n=5, Mann-Whitney U test. **(F)** Protein level of SASP factors: IL-1β, IL-6 and sICAM-1 in young donor-derived HAECs transfected with pre-miR-34a or negative control measured by ELISA. n=4, Mann-Whitney U test. **(G)** Expression of miR-34a-5p in young donor-derived HAECs transfected with si*NFE2L2*, si*TP53* or negative control. Relative expression was measured by real-time PCR. U6 snRNA served as an internal control. n=5, two-way ANOVA + Tukey’s. **(H)** Percentage of cells with increased SA-β-gal activity in young donor-derived HAECs transfected with si*NFE2L2*, anti-miR-34a-5p or negative control. n=5, two-way ANOVA + Tukey’s.

The expression of miR-34a was also elevated in young NRF2-deficient HAECs (Fig. 1G), which undergo premature senescence.^12^ This elevation was p53-dependent, as it was completely abrogated by p53 knock-down (Fig. 1G). Moreover, the increased expression of miR-34a led to premature senescence, since concomitant silencing of NRF2 and miR-34a abrogated the effect of NRF2 deficiency (Fig. 1H). Thus, we demonstrated a strong association of miR-34a with physiological and NRF2-dependent premature ageing of human aortic endothelial cells and critical role of p53 in this process.

### miR-34a governs the intima senescent phenotype in NRF2 tKO mice

Our previous study showed that disruption of NRF2 signalling drives the premature senescence of the murine aorta. Increased SA-β-gal activity can be found in all three layers of the aorta of young NRF2 tKO mice.^13^ Given that the NRF2-deficient HAECs are prematurely senescent and express miR-34a at high levels (Fig. 1G, H), we analysed the expression of miR-34a in the intima layer of NRF2 tKO aorta. *In situ* miR-34a hybridisation in *en face* projection of the aorta showed a strong elevation of the miR-34a levels in the EC layer compared to WT counterparts (Fig. 2A).

**Fig. 2.**
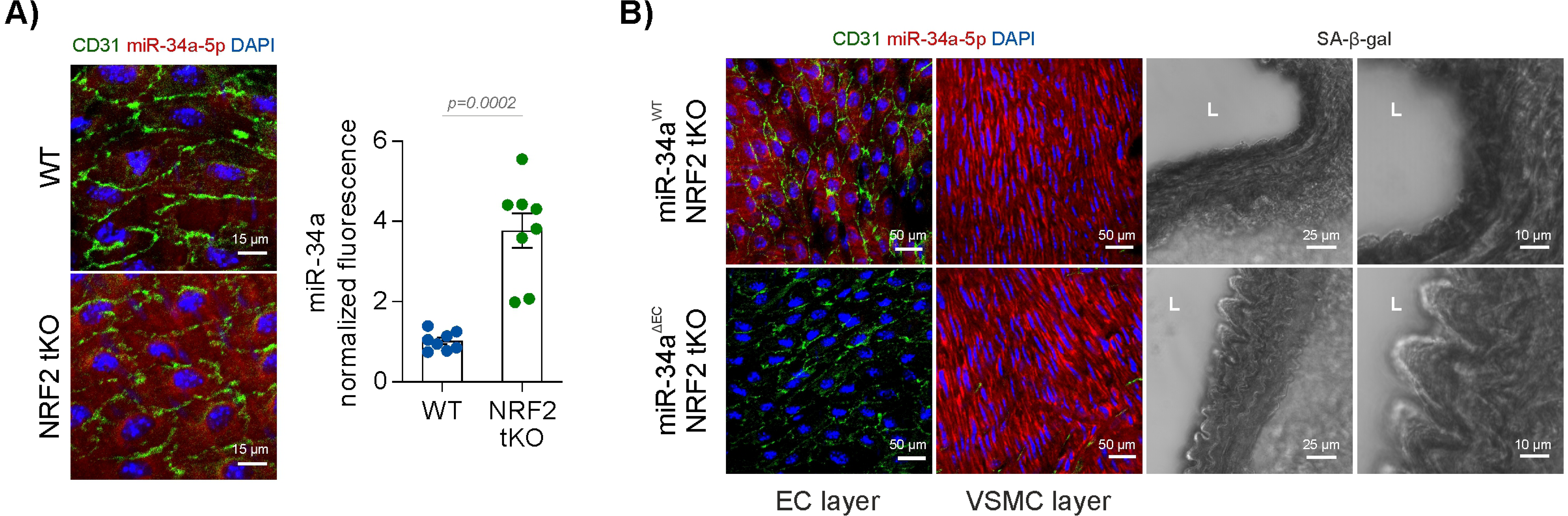
miR-34a level in the intimal layer of the murine aorta. **(A)** Expression of miR-34a-5p in the endothelial cells of the aortas from WT and NRF2 tKO mice measured by fluorescent *in situ* hybridization in *en face* projection. Representative pictures and quantitative data. 3-month-old mice, n=8, Mann-Whitney U test. Scale bar 15 µm. **(B)** Expression of miR-34a-5p in endothelial cells (EC) and vascular smooth muscle cells (VSMC) of the aortas from miR-34a^WT^/NRF2 tKO and miR-34a^ΔEC^/NRF2 tKO mice measured by fluorescent *in situ* hybridization in *en face* projection. Assessment of SA-β-gal activity, shown in black colour, in the cross section of the aortas from miR-34a^WT^/NRF2 tKO and miR-34a^ΔEC^/NRF2 tKO mice. 3-month-old mice, n=5-6. Representative pictures. L – lumen. Scale bar 50 µm for immunofluorescence; 25 µm and 10 µm for SA-β-gal activity, left and right panel, respectively.

To examine whether the senescent phenotype of NRF2 tKO aortas is driven by miR-34a, we generated a triple mutant model by crossing NRF2 tKO mice with inducible EC-specific miR-34a knockouts (miR-34a floxed mice crossed with tamoxifen-induced VE-cadherin-driven Cre recombinase-bearing strain). It allowed the specific deletion of miR-34a in endothelial cells, whereas its levels in vascular smooth muscle cells (VSMCs) remained unchanged (Fig. 2B). As shown in Fig. 2B, specific deletion of miR-34a in the endothelial layer of the NRF2 tKO aorta reduced the activity of SA-β-gal selectively in ECs but not in the media or adventitia. These results are in line with the data obtained in HAECs and demonstrate that premature senescence of mouse and human ECs driven by deficient NRF2 signalling depends on miR-34a.

### The knockout of endothelial miR-34a protects against AAA development

To address the role of endothelial miR-34a in AAA formation, we used the angiotensin II (Ang II) exposure model. In the first step, we examined how Ang II influences miR-34a levels in HAECs, mouse serum and mouse aortic wall. Treatment of HAECs with Ang II increased the expression of miR-34a (Fig. 3A). Alike, elevated levels of miR-34a were seen is sera of WT mice infused with Ang II. Remarkably, the highest levels of miR-34a, reaching 4-5 fold increase, were found in sera of animals which developed AAA (Fig. 3B). Moreover, infusion of Ang II upregulated miR-34a in the aortic wall in AAA forming mice, selectively in the layer of endothelial cells (Fig. 3C).

**Fig. 3.**
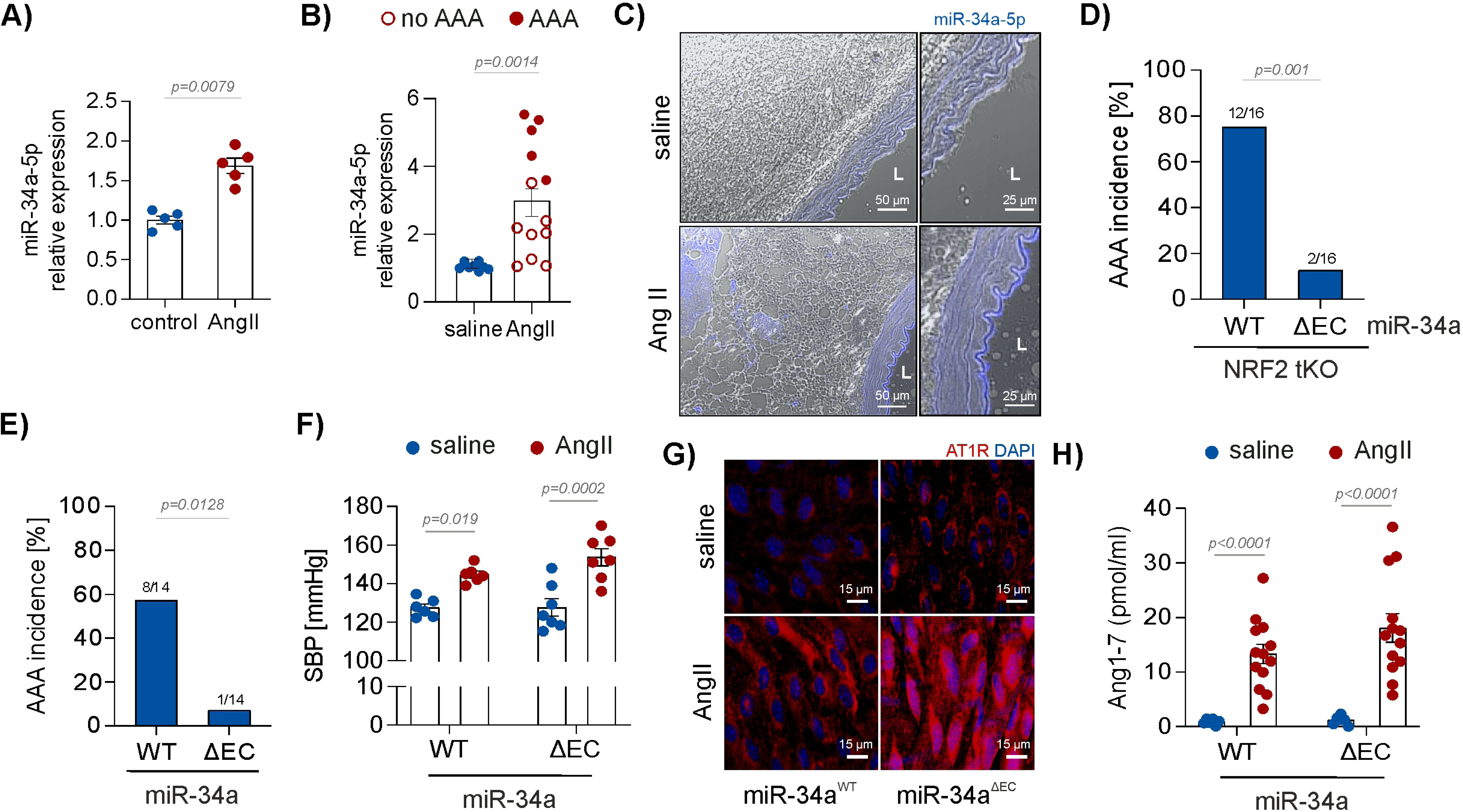
Effect of miR-34a endothelial deletion on AAA formation. **(A)** Expression of miR-34a-5p in young donor-derived HAECs stimulated with 500 nM Ang II for 6 h. Relative expression was measured by real-time PCR. U6 snRNA served as an internal control. n=5, Mann-Whitney U test. **(B)** Expression of miR-34a-5p in the sera of mice administered with Ang II (1000 ng/min/kg) for 7 days. Relative expression was measured by real-time PCR. U6 snRNA served as an internal control. 3-month-old mice, n=8-13, Mann-Whitney U test. **(C)** Representative images of conventional RNA *in situ* hybridization-based detection of miRNA-34a-5p in control and Ang II (1000 ng/min/kg, for 7 days) receiving animals. 3-month-old mice, n=5-8, L - lumen. Scale bar 50 µm and 25 µm for the left and right panel, respectively. **(D)** Incidence of AAA in miR-34a^WT^/NRF2 tKO and miR-34a^ΔEC^/NRF2 tKO mice administered with Ang II (1000 ng/min/kg) for 14 days. 3-month-old mice. n=16, Fisher’s exact test. **(E)** Incidence of AAA in miR-34a^WT^ and miR-34a^ΔEC^ mice administered with Ang II (1000 ng/min/kg) for 14 days. 6-month-old mice, n=14, Fisher’s exact test. **(F)** Systolic blood pressure in miR-34a^WT^ and miR-34a^ΔEC^ mice administered with Ang II (1000 ng/min/kg) for 3 days measured by tail-cuff plethysmography. 6-month-old mice, n=6-7, two-way ANOVA + Tukey’s. **(G)** AT1R level in endothelial cells of the aorta harvested from miR-34a^WT^ and miR-34a^ΔEC^ mice administered with Ang II (1000 ng/min/kg) for 3 days. Representative pictures of the *en face* immunofluorescent staining. 6-month-old mice, n=7. Scale bar 15 µm. **(H)** Ang1-7 level in the sera of miR-34a^WT^ and miR-34a^ΔEC^ mice administered with Ang II (1000 ng/min/kg) for 3 days, measured by ELISA. 6-month-old mice, n=6-14, two-way ANOVA + Tukey’s.

Taking into account the high predisposition of young NRF2 tKO mice to AAA development,^5^ elevated expression of miR-34a in aortic ECs of NRF2 tKO (Fig. 2A), and in response to the Ang II infusion seen solely in the intima layer of the aorta (Fig. 3C), in the next step we examined the AAA development rate in young NRF2 tKO mice devoid of miR-34a in the EC. Endothelial deletion of miR-34a reduced the incidence of AAA from 75% to 12.5% in young NRF2 tKO mice (Fig. 3D). This shows that formation of AAA in the NRF2 tKO mice depends on endothelial miR-34a, and the expression of miR-34a is a harmful factor.

AAA development was also abrogated in 6-month-old miR-34a^ΔEC^ mice with normal levels of NRF2 (Fig. 3E). The incidence of AAA was low in 3-month-old animals regardless of the miR-34a genotype (4 out of 24 WT mice and 5 out of 29 miR-34a^ΔEC^ mice). The protective effect of endothelial miR-34a deletion in 6-month old mice was not related to impaired signalling from Ang II, since *i)* Ang II induced hypertension in a similar manner in both experimental groups (Fig. 3F), *ii)* the level of the Ang II receptor, AT1R, was the highest in miR-34a^ΔEC^ aorta (Fig. 3G), and *iii)* concentrations of the vasodilative derivative of Ang II, angiotensin (1-7), were comparable in both groups (Fig. 3H). Therefore, it is rather doubtful that the alterations in the Ang II signalling account for the abrogation of AAA formation in miR-34a^ΔEC^ mice.

### miR-34a^ΔEC^ mice show a robust dysfunction-like phenotype of ECs

Healthy endothelial phenotype is compromised in the presence of cardiovascular risk factors and endothelial cell dysfunction (ECD) interrelates with cardiovascular diseases.^27^ Experimental studies showed that AAA formation is aggravated in the absence of endothelial nitric oxide synthase (eNOS),^28^ the enzyme producing NO, which is a sentinel of healthy endothelium and blood vessel homeostasis. In line, clinical data reported an inverse correlation between the maximum diameter of the aneurysm and flow-mediated dilation (FMD) in AAA patients.^29,30^ Therefore, having observed the protective effect of miR-34 endothelial deficiency on AAA development, we expected the improved or at least undisturbed endothelial cell function in miR-34a^ΔEC^ mice.

To investigate the functional status of the endothelium and blood vessels in these animals, we analysed several age-related features of endothelial dysfunction, namely quality of endothelial glycocalyx, stiffness of endothelial cells and aorta, production of NO and anti-inflammatory potential. Since we observed the formation of AAA rescued by endothelial miR-34a deletion in the 6-month-old but not 3-month old mice, functional experiments were carried out in mice of different age to better discriminate the potential influence of endothelial (dys)function on AAA development.

Taking the first step, we inspected the glycocalyx length and density and endothelial cell stiffness by atomic force microscopy (AFM). It revealed that miR-34a endothelial deficiency led to a decrease in glycocalyx coverage both in 3- and 6-month-old mice (Fig. 4A). The glycocalyx was shorter in older WT animals than in young WT mice, and in both younger and older miR-34a^ΔEC^ mice (Fig. 4B). Abnormal glycocalyx composition was also confirmed in mice deficient in endothelial miR-34a by immunofluorescent staining of N-acetyl-D-glucosamine and sialic acid components detected with WGA lectin (Fig. 4C). These data indicate aberrant glycocalyx structure and glycocalyx damage in miR-34a^ΔEC^ mice.

**Fig. 4.**
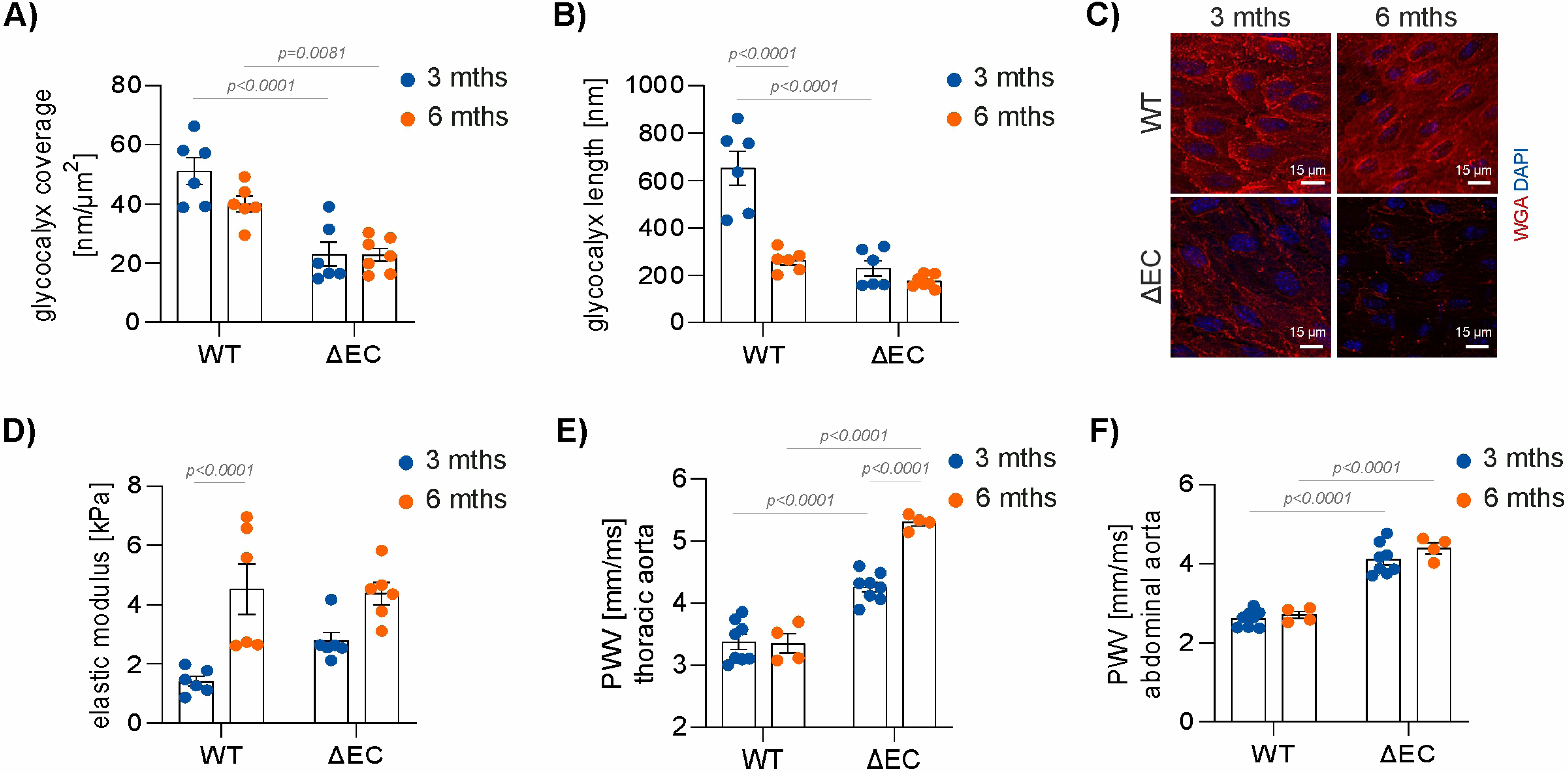
Assessment of endothelial dysfunction in miR-34a^ΔEC^ mice. **(A-B)** Measurement of endothelial cell glycocalyx (A) coverage and (B) length in the aorta of miR-34a^WT^ and miR-34a^ΔEC^ mice by atomic force microscopy. 3- and 6-month-old mice, n=6-7, two-way ANOVA + Tukey’s. **(C)** Representative images of the staining of glycocalyx components by Wheat Germ Agglutinin (WGA). 3- and 6-month-old mice, n=5. Scale bar 15 µm. **(D)** Measurement of the aortic endothelial cell stiffness in miR-34a^WT^ and miR-34a^ΔEC^ mice by atomic force microscopy. 3- and 6-month-old mice, n=6-7, two-way ANOVA + Tukey’s. **(E-F)** Pulse wave velocity (PWV) in (E) thoracic and (F) abdominal aorta was assessed by Doppler Flowmetry. 3- and 6-month-old mice, n=4-8, two-way ANOVA + Tukey’s.

Using AFM, we also observed that ageing significantly elevated aortic wall stiffness at the endothelial layer in WT mice, and there was a clear tendency for the increased stiffness in miR-34a^ΔEC^ aortas of young and older animals (Fig. 4D). Artery stiffening in the absence of endothelial miR-34a was also confirmed by the pulse wave velocity measurement in thoracic and abdominal aorta (Fig. 4E,F). Ageing aggravated this miR-34a-dependent effect in the thoracic part of the aorta (Fig. 4E).

Then we measured nitric oxide production in the aorta using electron paramagnetic resonance (EPR) with diethyldithiocarbamic acid sodium salt (DETC) as a spin-trap. Production of NO was compromised in 3-month-old miR-34a^ΔEC^ mice (7.98±0.48 vs 5.52±0.47 NO-DETC signal per µg of aorta tissue for WT and miR-34a^ΔEC^ mice, respectively; p=0.0029). In 6-month-old animals production of NO was below the threshold of EPR detection, regardless the mir-34a status.

In the last step, we used the intravital microscopy, which permitted life imaging of TNFα-induced leukocyte adhesion to mesenteric venules, which reflects the inflammatory potential of the vessel intima. Surprisingly, also here the removal of endothelial miR-34a, which prevents AAA development, was associated with the abnormal phenotype of miR-34a^ΔEC^ vascular bed. We found a significantly increased number of cells adhering to the vessel wall in miR-34a^ΔEC^ mice and a notable decrease in leukocyte velocity irrespectively of the age of the mice (Fig. 5A-C).

**Fig. 5.**
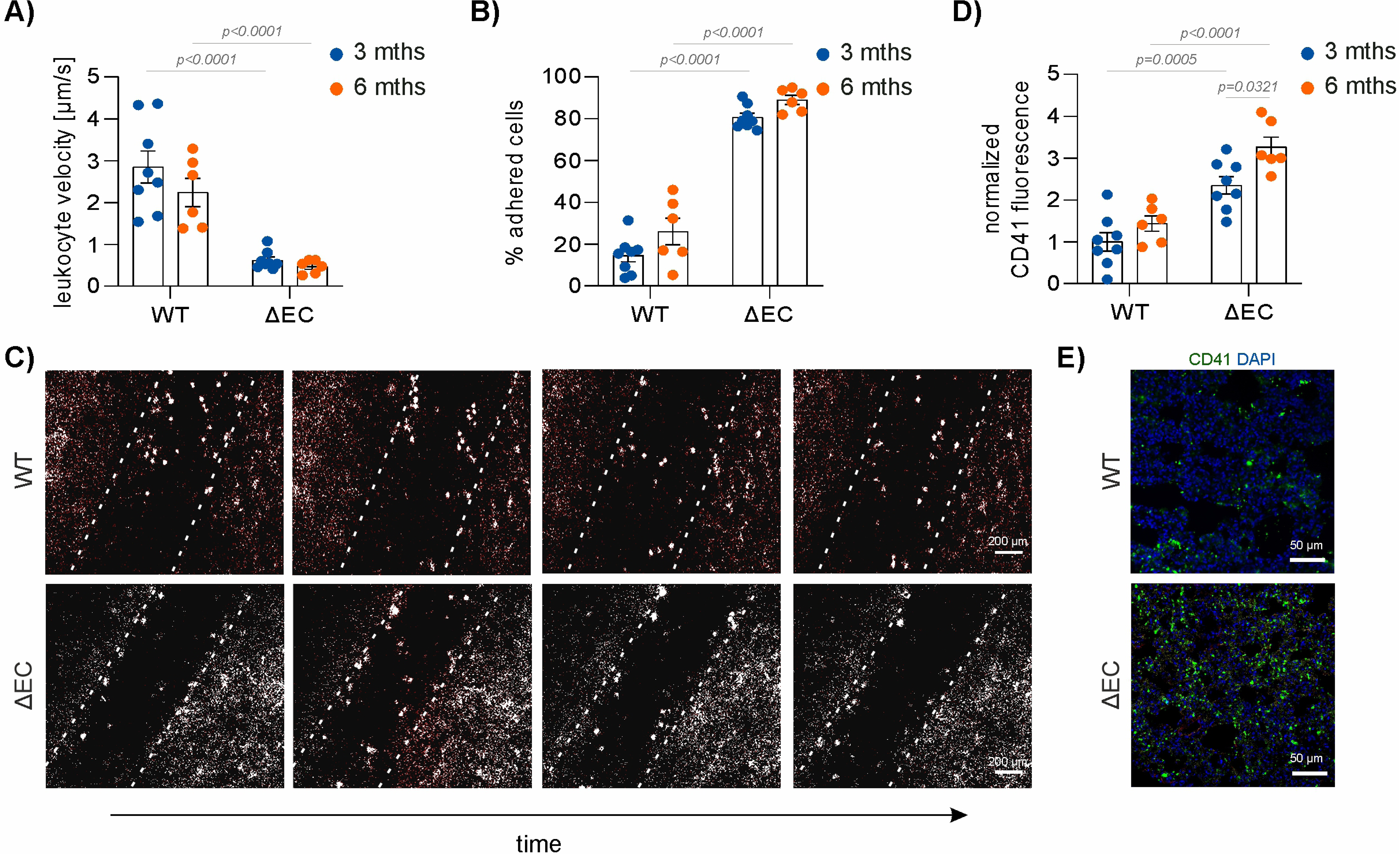
Assessment of proinflammatory and prothrombotic phenotype of blood vessels in miR-34a^ΔEC^ mice. **(A-C)** Measurement of (A) the leukocyte velocity and (B) percentage of adhered leukocytes in the mesenteric venules of miR-34a^WT^ and miR-34a^ΔEC^ mice by intravital microscopy. (C) Representative pictures of adhered leukocytes. 3- and 6-month-old mice, n=6-8, two-way ANOVA + Tukey’s. Scale bar 200 µm. **(D-E)** Immunofluorescent staining of platelets in the lungs of miR-34a^WT^ and miR-34a^ΔEC^ mice. Collagen-epinephrine model of thrombosis. (D) Quantitative data and (E) representative pictures. Anti-CD41 antibody was used. 3- and 6-month-old mice, n=6-8, two-way ANOVA + Tukey’s. Scale bar 50 µm.

Furthermore, the grade of collagen/epinephrine-induced pulmonary thrombosis, measured by CD41 immunostaining in lungs, was elevated in mice devoid of endothelial miR-34a, although without affecting the platelet coagulation features (Fig. S1). In the case of pulmonary thrombosis, the effect was stronger in 6-month-old mice, compared to younger animals (Fig. 5D,E). To conclude, our results unexpectedly indicate that removal of miR-34a from endothelial cells protects against AAA formation, but at the same time exacerbates features typical of endothelial dysfunction. No analysed parameter influenced by miR-34a deletion, reflected the risk of AAA development in animals of different ages. Therefore, we suppose that ECD is not a predominant mechanism responsible for AAA formation, at least in this experimental setting.

### High intimal EC number is found in NRF2 tKO miR-34a^ΔEC^ and miR-34a^ΔEC^ aortas, which are protected from AAA

Since the protective effect of endothelial miR-34a deficiency on AAA formation cannot be attributed to the maintenance of endothelial cell function or alterations in Ang II signalling, seeking for a possible mechanism, we inspected the luminal surface of the NRF2 tKO miR-34a^ΔEC^ and miR-34a^ΔEC^ aortas of Ang II-infused mice, in relation to respective control groups. Interestingly, we found a higher number of intimal ECs in Ang II-treated mice devoid of endothelial miR-34a (Fig. 6A,B).

**Fig. 6.**
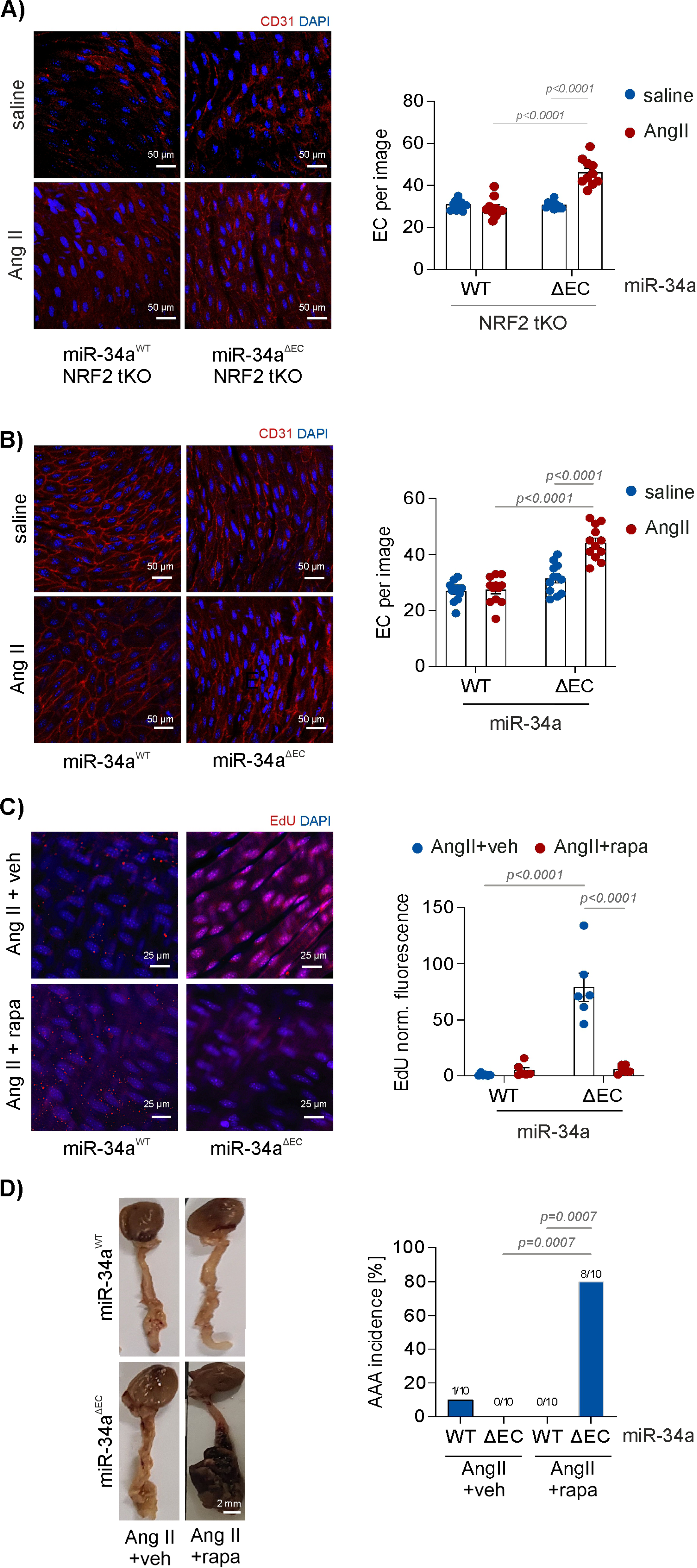
Effect of intimal cell proliferation on AAA formation in mice. **(A)** Immunofluorescent *en face* aorta staining for intimal ECs in miR-34a^WT^/NRF2 tKO and miR-34a^ΔEC^/NRF2 tKO mice administered with Ang II (1000 ng/min/kg) for 3 days. Representative pictures and quantitative data. 3-month-old mice, n=12-15, two-way ANOVA + Tukey’s. Scale bar 50 µm. **(B)** Immunofluorescent *en face* aorta staining for intimal ECs in miR-34a^WT^ and miR-34a^ΔEC^ mice administered with Ang II (1000 ng/min/kg) for 3 days. Representative pictures and quantitative data. 6-month-old mice, n=12-15, two-way ANOVA + Tukey’s. Scale bar 50 µm. **(C)** Immunofluorescent *en face* aorta staining for proliferating intimal ECs in miR-34a^WT^ and miR-34a^ΔEC^ mice administered with Ang II (1000 ng/min/kg) and rapamycin (4 mg/kg) for 8 days. Representative pictures and quantitative data. 3-month-old mice, n=6, two-way ANOVA + Tukey’s. Scale bar 25 µm. **(D)** Incidence of AAA in miR-34a^WT^ and miR-34a^ΔEC^ mice administered with Ang II (1000 ng/min/kg) and rapamycin (4 mg/kg) for 7 days. Representative pictures of the aortas and quantitative data. 3-month-old mice, n=10, Fisher’s exact test. Scale bar 2 mm.

Therefore, in the next step, we aimed to block endothelial proliferation to verify if it influences AAA development in miR-34a^ΔEC^ mice. We injected miR-34a^WT^ and miR-34a^ΔEC^ mice with rapamycin, which is known to hamper the proliferation of ECs.^31^ Indeed, rapamycin administration efficiently decreased intimal cell proliferation (Fig. 6C, S2) and simultaneously led to robust induction of AAA only in miR-34a^ΔEC^ (Fig. 6D). Moreover, macroscopically, the formed AAA seemed rupture-prone (Fig. 6D). Therefore, we conclude that miR-34a-regulated increase in EC proliferation inversely correlates with the AAA susceptibility. Consequently, it can be supposed that EC proliferation could confer protection against AAA.

### MTA2 is the target of miR-34a that regulates Ang II-induced EC proliferation

To identify a target of miR-34a mediating the Ang II-induced proliferative effect of miR-34a deficiency, we used age donor-derived HAECs. A canonical target of miR-34a is sirtuin 1 (SIRT1), and the protective impact of miR-34a deficiency is usually attributed to the increase of this protein.^32^ However, concomitant downregulation of SIRT1 and miR-34a (Fig. S3A, B) in aged-donor derived HAECs treated with Ang II did not reverse the high proliferation capacity of the cells (Fig. S3C). Thus, SIRT1 is not a mediator of miR-34a-dependent regulation of proliferation.

Therefore, in the next step, we performed mass spectrometry (MS) analysis to identify other potential proteins that could mediate miR-34a-dependent effects in endothelial cells. It revealed 54 proteins differentially expressed between HAECs derived from aged and young donors (Fig. 7A, Tab. S1), with the highest statistical score (p=0.002) and the fold change 0.24 aged-donor vs young donor for metastasis-associated protein 2 (MTA2), one of the miR-34a targets,^33^ deacetylating p53.^34^ The following MS analysis of aged donor HAECs transfected with anti-miR-34a or control sequence showed 41 significantly changed proteins (Fig. 7B, Tab. S2). Among them, 29 proteins were reported to regulate cell proliferation with positive or negative impact (Fig. 7C), and MTA2 was one of them. To assess the proliferation status of anti-miR-34a-transfected HAECs compared to control cells, we performed PCNA staining that showed increased number of PCNA-positive cells devoid of miR-34a for young- and aged-donor derived cells (Fig. 7D). The percentage of PCNA positive cells was further increased in anti-miR-34a HAECs exposed to Ang II (Fig. 7E), which is in line with the *in vivo* observations (Fig. 6). Interestingly, MTA2 level was elevated 3.47 times (p=0.029) in HAECs deficient in miR-34a compared to control counterparts (Fig. 7B, Tab. S2). The increase in MTA2 in miR-34a knockdown HAECs, strengthened by Ang II treatment, was also confirmed by immunofluorescence staining (Fig. 7F).

**Fig. 7.**
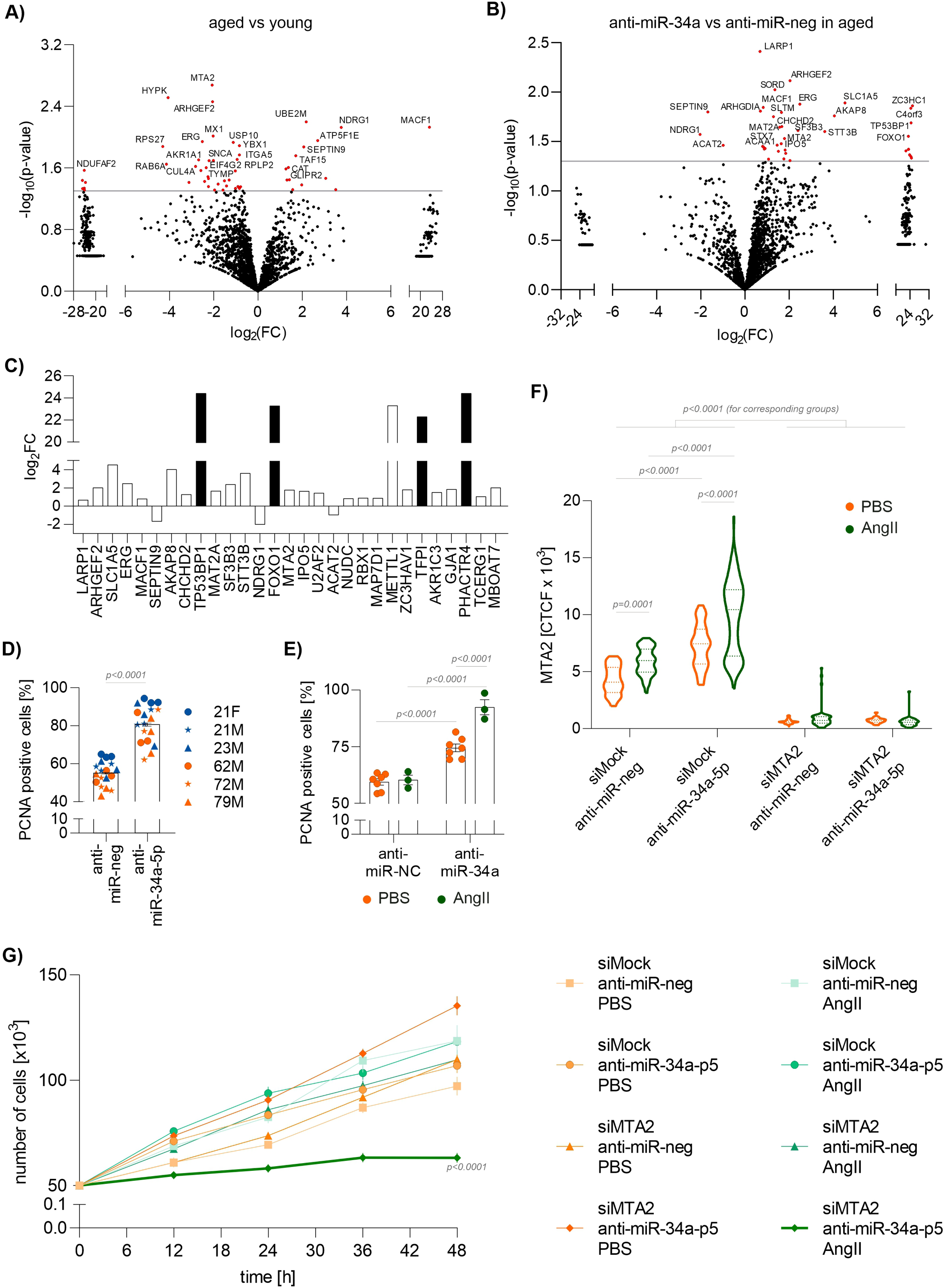
Effect of MTA2 on EC proliferation induced by Ang II. **(A)** Proteome profiling in young and aged donor-derived HAECs. Whole-cell lysates were analysed by mass spectrometry. FC – fold change, n=9, Mann-Whitney U test. **(B)** Proteome profiling in aged donor-derived HAECs transfected with anti-miR-34a or scrambled sequence. Whole-cell lysates were analysed by mass spectrometry. FC – fold change, n=9, Mann-Whitney U test. **(C)** Proliferation-related proteins with a significantly different level in anti-miR-34a aged donor-derived HAECs in comparison to anti-miR-neg transfected cells. Whole-cell lysates were analysed by mass spectrometry. FC – fold change, n=9. White and black bars – positive and negative regulators of proliferation, respectively. **(D)** PCNA immunofluorescent staining of young and aged donor-derived HAECs transfected with anti-miR-34a or scrambled sequence. Quantitative data, percentage of positive cells. n=18, Mann-Whitney U test. **(E)** PCNA immunofluorescent staining of aged donor-derived HAECs transfected with anti-miR-34a or scrambled sequence and stimulated with Ang II (500 nM) for 6 h. Quantitative data, percentage of positive cells. n=3-7, two-way ANOVA + Tukey’s. **(F)** MTA2 immunofluorescent staining of aged donor-derived HAECs transfected with anti-miR-34a, si*MTA2* or scrambled sequence and stimulated with Ang II (500 nM) for 6 h. Quantitative data, CTCF - corrected total cell fluorescence. n=28-84, two-way ANOVA + Tukey’s. **(G)** Growth curve of aged donor-derived HAECs transfected with anti-miR-34a, si*MTA2* or scrambled sequence and stimulated with Ang II (500 nM). Ang II was added at time point 0 h. n=6, two-way ANOVA + Tukey’s.

Finally, we examined whether downregulation of MTA2 inhibits the Ang II-stimulated proliferation of miR-34a-deficient HAECs. The concomitant silencing of miR-34a and MTA2 led to the complete abrogation of HAEC cell growth. The effect was visible only in cells exposed to Ang II (Fig. 7G). To check if this effect is specific, we conducted a second set of experiments in which anti-miR-34a/siMTA2 cells were treated with Ang II with 24 h delay. The addition of Ang II to media ceased the growth of these cells, but not the respective control (Fig. S4). In conclusion, miR-34a orchestrates Ang II-induced HAECs proliferation in MTA2-dependent manner.

## Discussion

In the present study, we reveal a clear disassociation between ECD and AAA susceptibility and show a predominant role for endothelial miR-34a deletion in protecting against AAA development. This implies that dysfunctional endothelium might be still protective, at least if it maintains proliferation.

It is well-established that AAA formation is accompanied by extracellular matrix degradation, immune cell infiltration, and SMC loss.^35^ However, none of these pathological alterations have been validated as a good drug target for effective therapy. This could be related to aiming at too late stages of AAA development.^1^ Another open concept of AAA pathomechanisms is based on ECD,^36^ which is commonly accepted to predispose to CVDs and to be central in cardiovascular system pathology.^27,37^ By definition, ECD is a spectrum of phenotypic states associated with impaired ability of endothelial cells to regulate vascular tone, maintain thromboresistance, prevent leukocyte adhesion, vessel wall inflammation, and permeability.^37^ It is proposed that loss of endothelial function is gradual and accumulation of dysfunctional phenotypes is linked to CVDs of varying severity.^37^

The critical finding of our study demonstrates, however, a clear disconnection between miR-34a-prevented ECD and formation of AAA, which does not fit to the current view of the importance of ECD in AAA pathogenesis.^36^ miR-34a^ΔEC^ mice show a robust dysfunction-like phenotype of endothelium in various vascular beds, including aorta, lung microvessels, and mesenteric vein, represented by degraded glycocalyx layer, increased adhesion of leukocytes and thrombosis, reduced NO production and vessel wall stiffening. On the other hand, they are fully protected against Ang II-induced AAA development, even on the AAA-prone NRF2 KO^5^ background. Therefore, we suppose that ECD associated with AAA, observed in AAA patients and in animal models,^36^ could be considered a response to the development of AAA rather than its cause.

This suggestion is corroborated by experimental data obtained from mice deficient in eNOS. Indeed, eNOS KO mice exhibit a strong ECD phenotype, including impaired acetylcholine-induced vasodilatation,^38^ increased leukocyte adhesion, platelet aggregation, and thrombosis; they are hypertensive and prone to stroke and atherosclerosis.^39^ However, to our knowledge, no reports show enhanced AAA formation in eNOS KO mice. Similarly, calcium chloride-induced aneurysms in carotid arteries are even smaller in these animals compared to their WT counterparts.^40^ The eNOS-dependent aggravation of AAA development was shown only in ApoE KO animals fed a Western-type diet,^28^ but in this case the experimental setting is strongly affected by the effects related to the ApoE null background. An additional supporting argument arises from clinical studies. Diabetes was shown to be negatively associated with AAA growth in patients,^41^ but is highly correlated with ECD.^42^

The important finding of our study points to endothelial miR-34a as imperative regulator of AAA development. Young NRF2 tKO miR-34a^ΔEC^ and 6-month old miR-34a^ΔEC^ mice challenged with Ang II exhibit strong intimal cell proliferation and are protected from AAA. This effect is reversed by rapamycin, which blocks EC division.^31^ Our results indicate that one of the potential mechanism conferring protection to the aortic wall could be EC proliferation, promoted by miR-34 deletion, which implies that maintaining a compact intimal barrier might be essential to prevent AAA.

Interestingly, telomeres are shortened in ECs present in human aneurysmal tissue,^43^ which can limit EC division. In line, the restoration of the endothelial lining by local endovascular infusion of ECs prevents AAA formation and stabilizes aneurysm expansion in a guinea pig/rat xenograft model of AAA.^44^ This concept is further supported by the recent paper showing that disruption of endothelial tight junctions is an early critical step in the progression of a β-aminopropionitrile (BAPN)-induced mouse model of thoracic aortic aneurysm and dissection.^45^ Furthermore, it was shown that the size of elastase-induced AAA formed in rats inversely correlates with EC proliferation.^46^ The variety of aneurysm models used in the papers discussed above indicates that the EC-mediated maintenance of the barrier of the vessel wall could be the universal preventive mechanism against aneurysm.

EC and SMC turnover is estimated to be 1-3% in arteries.^47–50^ Single-cell transcriptional profiling of the aortic endothelium shows that more than 95% of cells exhibit a quiescent phenotype.^51^ ECs, however, may generate a robust proliferative response in injured or injury-prone areas to compensate for the cell loss.^49,52,53^ Also Ang II can induce aortic EC proliferation as we show in this study and which is consistent with the recently published paper.^50^ This replicative capacity of aortic ECs is compromised in ageing, and the ATF3 transcription factor was identified as the crucial regulator of this process.^52^

Our data show that the NRF2/miR-34a axis is also a critical modulator of the EC proliferative capacity in ageing. We recently reported that aortic ECs derived from aged donors express NRF2 at extremely low levels.^11^ Aortas of mice lacking NRF2 transcriptional activity are prematurely senescent^13^ and so are HAECs deficient in NRF2.^12^ This phenotype is associated with impaired EC proliferation,^13^ and can be counteracted by miR-34a knockdown, as we show in the present study.

The mechanism of miR-34a-dependent regulation of EC proliferation becomes particularly important in the context of protection against AAA. Endothelial deletion of miR-34a in NRF2 tKO mice counteracts the senescent phenotype of ECs, and unleashes the replicative capacity of ECs upon Ang II infusion. As our rescue experiments show, this effect could be attributed to the upregulation of MTA2 in ECs deficient in miR-34a, but not to the canonical miR-34a target, SIRT1. The other potential mediator of miR-34a-regulated EC proliferation arising from the mass spectrometry study could be ERG (ETS Transcription Factor ERG, p55) which is not, in contrast to MTA2,^33^ a direct target for miR-34a (mirdb.org). ERG controls EC proliferation and quiescence.^54,55^ Its level is significantly reduced in ageing HAECs (fold change 0.17 aged vs young donor-derived cells, p=0.011), which is reversed by miR-34a silencing (fold change 5.6 aged vs young donor-derived cells, p=0.013). However, we have not confirmed its significance in Ang II-induced miR-34a knockdown-mediated EC proliferation experimentally.

Interestingly, the control of EC proliferation and modulation of mechanisms that maintain healthy endothelial phenotype are uncoupled. Our data shows that ECs devoid of miR-34a display a dysfunctional-like phenotype, and at the same time, a similar phenotype is triggered by increased expression of miR-34a.^32^ Therefore, the fine-tuning of miR-34a level seems to be crucial for the maintenance of EC functional balance. However, EC proliferation is regulated in a single mode. In contrast to the silencing of miR-34a, its upregulation ceases the EC division.^56^

The clinical AAA scoring system requires the establishment of circulating biomarkers that would facilitate the evaluation of aneurysm development, progression and the risk of its rupture.^57^ Currently, the most commonly used clinical biomarker for AAA is D-dimer, which is related to the presence and size of intraluminal thrombus, the diameter and growth of AAA, but is not specific.^58^ There are also numerous other markers, that could be potentially used, such as fibrinogen, tissue plasminogen activator, PAI activity, metalloproteinases and tissue inhibitors of metalloproteinases, and others.^57^ Still, their clinical utility is limited, mostly due to the lack of disease specificity and the fact that AAA is a multifactorial disorder, so probably a panel of circulating biomarkers should be considered to achieve the adequate precision. Our experimental study shows that miR-34a is significantly elevated in sera of mice treated with Ang II, but the exceptionally high levels are found in sera of mice that develop AAA. Interestingly, elevated levels of miR-34a are also found in PBMCs from AAA patients,^21^ which may lay the grounds for large, prospective clinical studies assessing the utility of miR-34a in the prediction and progression of AAA.

There are two potential limitations of our study. In our experimental setting, rapamycin was administered systemically. Therefore, we cannot exclude its potential effects on other cell types, for example, SMCs. Although rapamycin blocks proliferation of EC efficiently, it can also affect other important biological processes, such as cellular metabolism. However, the additional experiments presented in this report strongly support the concept of EC proliferation-dependent protection against AAA. The other potential limitation is that all experiments concerning the status of endothelial dys(function) in miR-34a^ΔEC^ mice were performed without Ang II challenge. It would be, however, unlikely that Ang II injection could reverse the dysfunctional phenotype of ECs.^59^

## Conclusions

miR-34a represents a multifunctional rheostat of endothelial phenotype. Upregulated endothelial miR-34a mediates NRF2 deficiency-induced endothelial senescence, whereas miR-34 endothelial knockout results in the impairment of NO-dependent function, the loss of glycocalyx, vascular stiffness, leads to the activation of pro-adhesive, pro-thrombotic endothelial responses as well as robust endothelial Ang II-induced proliferation. Despite the detrimental effects of miR-34 deficiency on endothelial function, it protects against the development of AAA, suggesting that endothelial miR-34-dependent mechanisms, likely regulation of EC proliferation, determines the outcome of AAA.

## Supplementary material

**Fig. S1.**
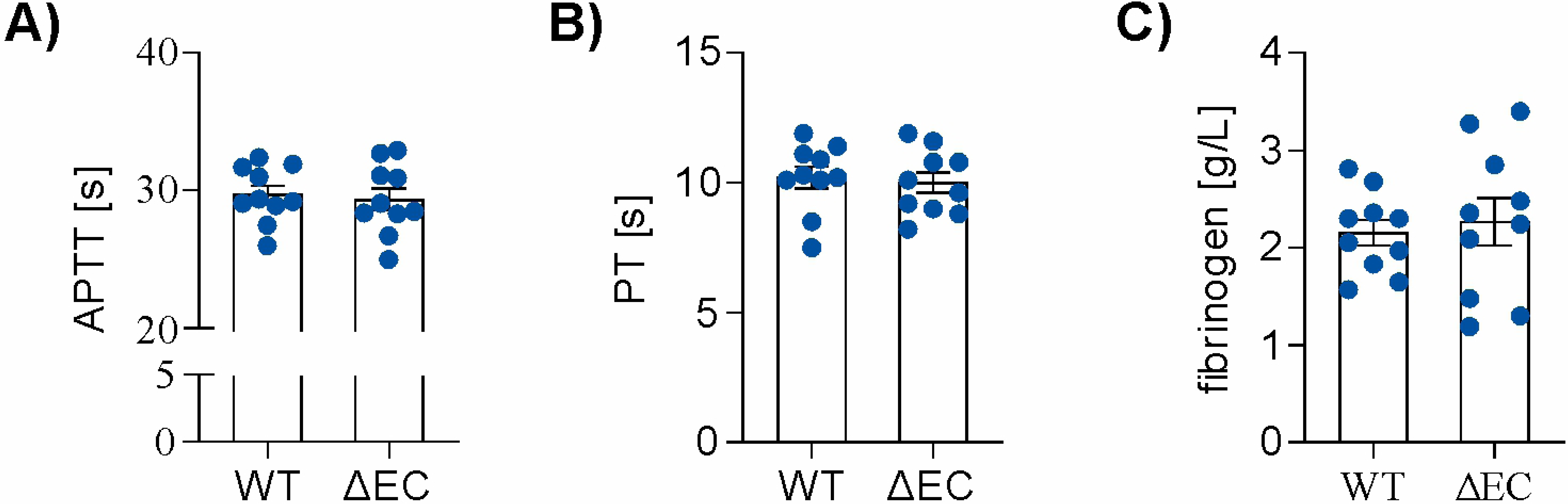
Effect of endothelial miR-34a deletion on the platelet coagulation parameters. **(A)** Activated partial thromboplastin time (APTT), **(B)** prothrombin time (PT) and **(C)** fibrinogen level in miR-34a^WT^ and miR-34a^ΔEC^ mice. 3-month-old mice, n=10, Mann-Whitney U test.

**Fig. S2.**
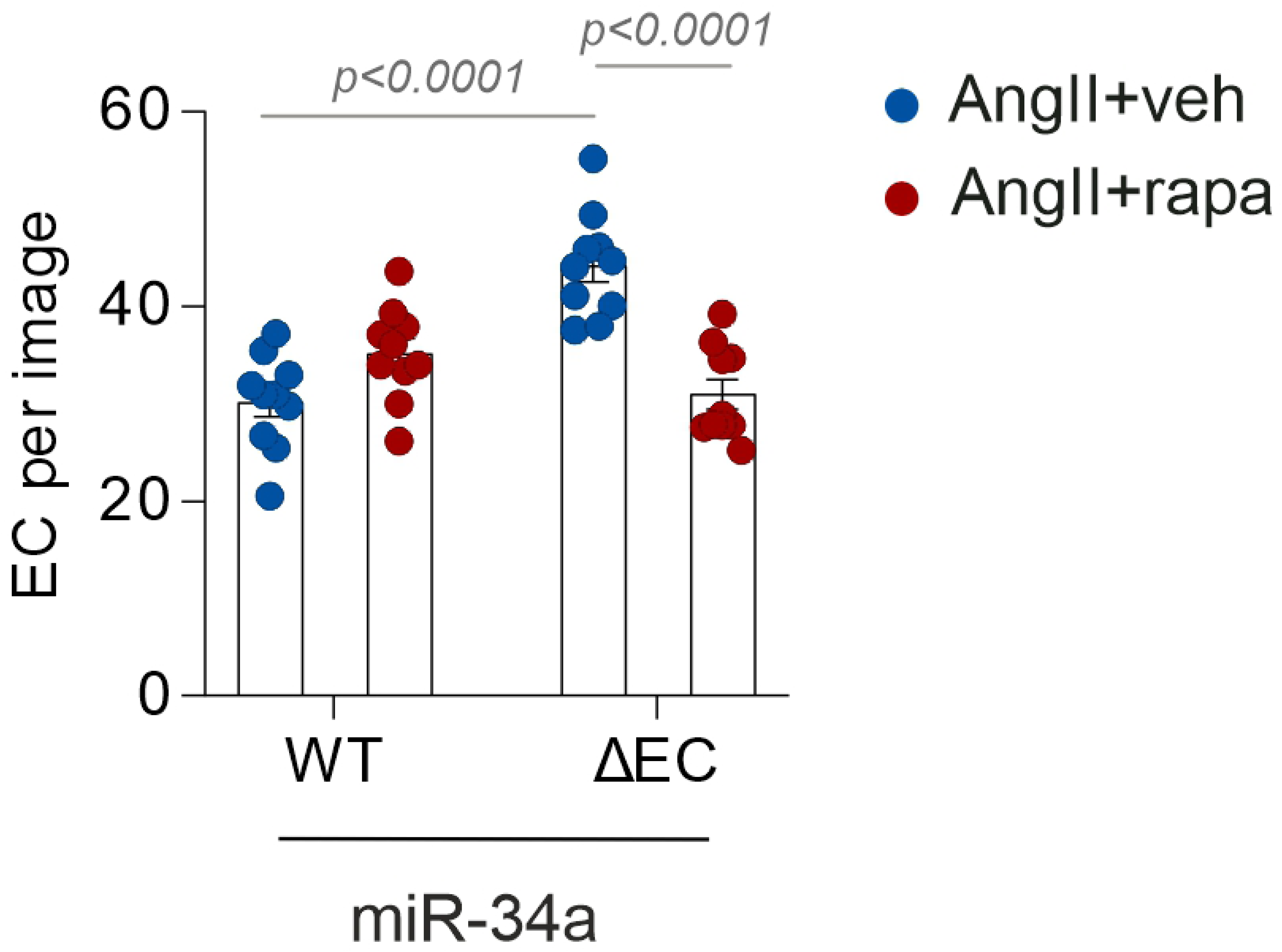
Effect of Ang II on EC proliferation in miR-34a^WT^ and miR-34a^ΔEC^ mice. Immunofluorescent *en face* aorta staining for proliferating intimal ECs in administered with Ang II (1000 ng/min/kg) and rapamycin (4 mg/kg) for 8 days. Quantitative data. 3-month-old mice, n=8-10, two-way ANOVA + Tukey’s.

**Fig. S3.**
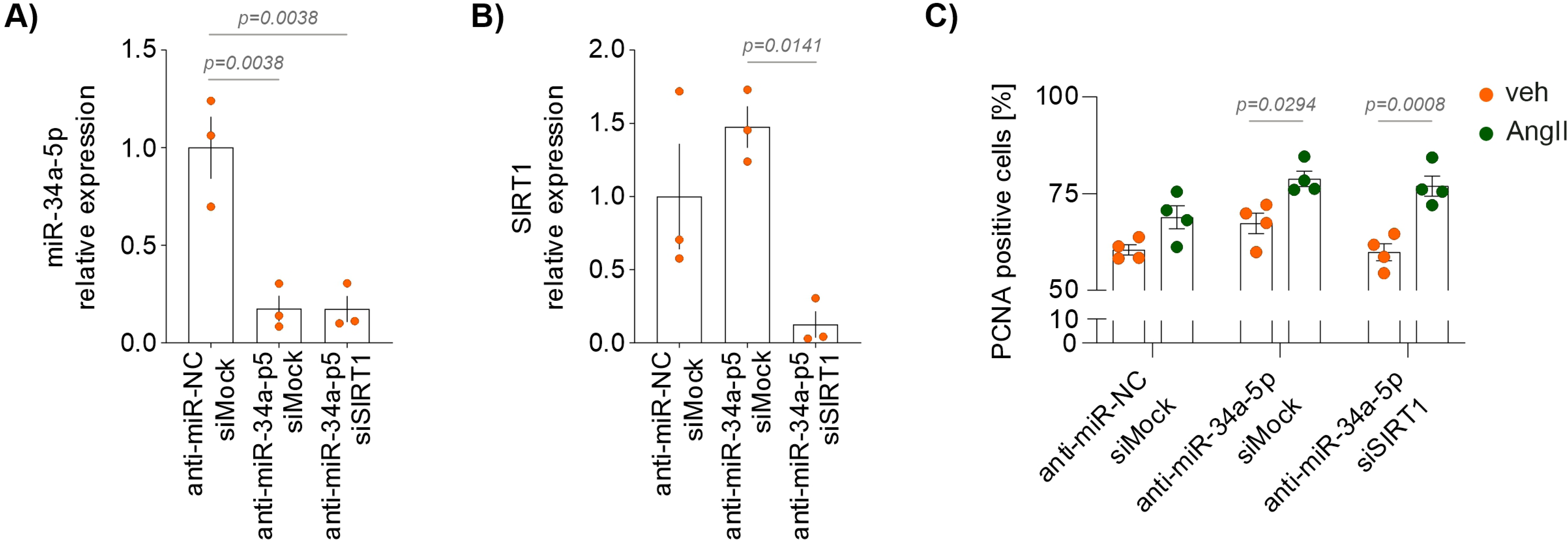
Effect of SIRT1 knockdown on HAEC proliferation. **(A)** Relative expression of miR-34a-p5 in aged donor-derived HAECs transfected with anti-miR-34a, si*SIRT1* or scrambled sequence. n=3, one-way ANOVA. **(B)** Relative expression of *SIRT1* in aged donor-derived HAECs transfected with anti-miR-34a, si*SIRT1* or scrambled sequence. n=3, one-way ANOVA. **(C)** PCNA immunofluorescent staining of aged donor-derived HAECs transfected with anti-miR-34a, si*SIRT1* or scrambled sequence and stimulated with Ang II (500 nM) for 6 h. Quantitative data, percentage of positive cells. n=4, two-way ANOVA + Tukey’s.

**Fig. S4.**
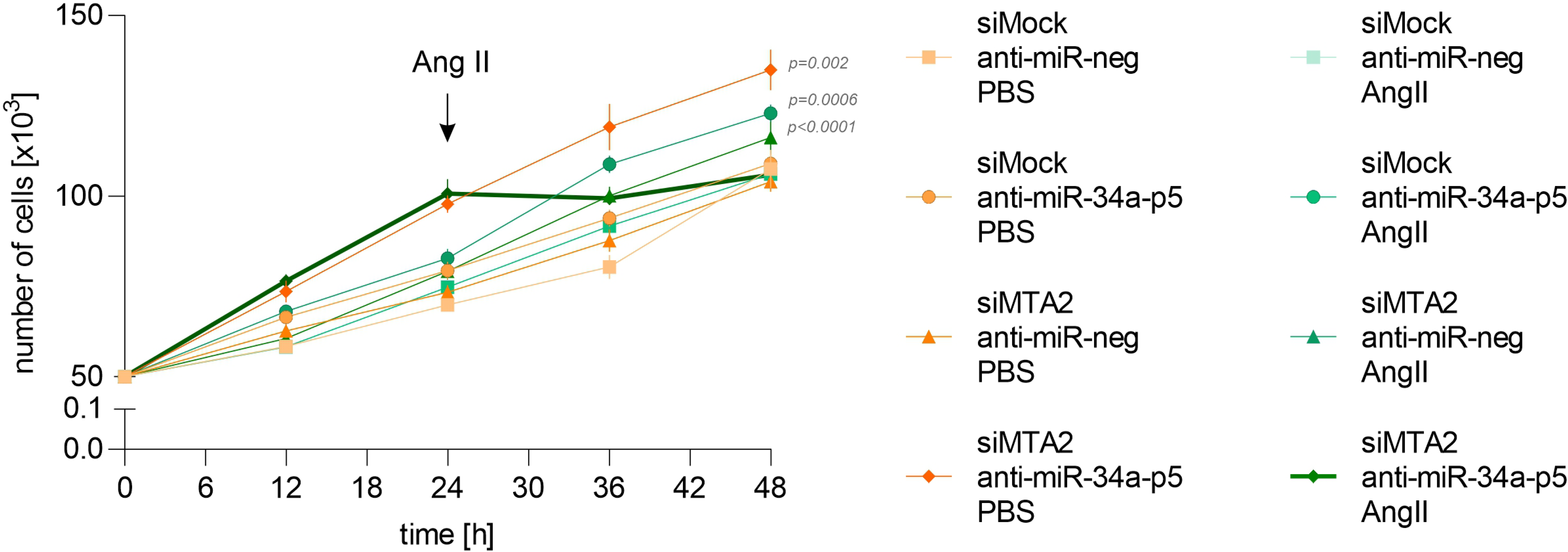
Effect of MTA2 on EC proliferation induced by Ang II. Growth curve of aged donor-derived HAECs transfected with anti-miR-34a, si*MTA2* or scrambled sequence and stimulated with Ang II (500 nM). Ang II was added at time point 24 h. n=6, two-way ANOVA + Tukey’s.

Tables S1, S2

## Acknowledgements

The authors acknowledge the staff at the Animal Facility for mice care and breeding and the administrative staff at the Department of Medical Biotechnology for their assistance. We thank Milena Cichon, Jakub Bartus and Beata Rysiewicz for technical help. Figures were created using BioRender.com.

## Authors’ contribution

Experiments were performed at the Faculty of Biochemistry, Biophysics and Biotechnology, Jagiellonian Centre for Experimental Therapeutics and Institute of Physics, Jagiellonian University. Proteome profiling was done at Mass Spectrometry Laboratory, Institute of Biochemistry and Biophysics, Polish Academy of Sciences. Conception and design of the work - AGP and AK; Funding acquisition - AGP, AB, AJ; The acquisition, analysis, and interpretation of data for the work - AK, DK, AB, MTK, DC, AGP; Writing - original draft, AGP and AK; Writing - review and editing, AK, DK, AB, MTK, DC, SC, AJ, AGP. All authors approved the final version of this manuscript and agreed to be accountable for all aspects of the work.

## List of abbreviations

AAA: abdominal aortic aneurysm
AAV: adeno-associated viral vectors
Ang II: angiotensin II
EC: endothelial cells
ECD: endothelial cells dysfunction
HAEC: human aortic endothelial cells
miR-34a^ΔEC^: deletion of miR-34a in endothelial cells
NO: nitric oxide
PAI: plasminogen activator inhibitor
PBMC: peripheral blood mononuclear cells
PWV: pulse wave velocity
SA-β-gal: senescence-associated β-galactosidase
SASP: senescence-associated secretory phenotype
SMC: smooth muscle cells
tKO: transcriptional knockout
WGA: wheat germ agglutinin
WT: wild type

## Data availability

The MS proteomics data have been deposited to the ProteomeXchange Consortium via the PRIDE partner repository^60^. The data underlying this article will be shared on reasonable request to the corresponding author.

## References

1. Golledge J. Abdominal aortic aneurysm: update on pathogenesis and medical treatments. Nat Rev Cardiol 2019;16:225–242.

2. Sakalihasan N, Michel J-B, Katsargyris A, Kuivaniemi H, Defraigne J-O, Nchimi A, Powell JT, Yoshimura K, Hultgren R. Abdominal aortic aneurysms. Nat Rev Dis Primers 2018;4:34.

3. Monteiro JP, Bennett M, Rodor J, Caudrillier A, Ulitsky I, Baker AH. Endothelial function and dysfunction in the cardiovascular system: the long non-coding road. Cardiovascular Research 2019;115:1692–1704.

4. Kloska D, Kopacz A, Piechota-Polanczyk A, Nowak WN, Dulak J, Jozkowicz A, Grochot-Przeczek A. Nrf2 in aging - Focus on the cardiovascular system. Vascul Pharmacol 2019;112:42–53.

5. Kopacz A, Werner E, Grochot-Przęczek A, Klóska D, Hajduk K, Neumayer C, Józkowicz A, Piechota-Polanczyk A. Simvastatin Attenuates Abdominal Aortic Aneurysm Formation Favoured by Lack of Nrf2 Transcriptional Activity. Oxid Med Cell Longev 2020;2020:6340190.

6. Baird L, Yamamoto M. The Molecular Mechanisms Regulating the KEAP1-NRF2 Pathway. Mol Cell Biol 2020;40:e00099–20.

7. Dinkova-Kostova AT, Copple IM. Advances and challenges in therapeutic targeting of NRF2. Trends Pharmacol Sci 2023;44:137–149.

8. Trougakos IP. Nrf2, stress and aging. Aging (Albany NY) 2019;11:5289–5291.

9. Ungvari Z, Bailey-Downs L, Sosnowska D, Gautam T, Koncz P, Losonczy G, Ballabh P, Cabo R de, Sonntag WE, Csiszar A. Vascular oxidative stress in aging: a homeostatic failure due to dysregulation of NRF2-mediated antioxidant response. Am J Physiol Heart Circ Physiol 2011;301:H363–372.

10. Ungvari Z, Bailey-Downs L, Gautam T, Sosnowska D, Wang M, Monticone RE, Telljohann R, Pinto JT, Cabo R de, Sonntag WE, Lakatta EG, Csiszar A. Age-associated vascular oxidative stress, Nrf2 dysfunction, and NF-{kappa}B activation in the nonhuman primate Macaca mulatta. J Gerontol A Biol Sci Med Sci 2011;66:866–875.

11. Kopacz A, Kloska D, Targosz-Korecka M, Zapotoczny B, Cysewski D, Personnic N, Werner E, Hajduk K, Jozkowicz A, Grochot-Przeczek A. Keap1 governs ageing-induced protein aggregation in endothelial cells. Redox Biol 2020;34:101572.

12. Kloska D, Kopacz A, Cysewski D, Aepfelbacher M, Dulak J, Jozkowicz A, Grochot-Przeczek A. Nrf2 Sequesters Keap1 Preventing Podosome Disassembly: A Quintessential Duet Moonlights in Endothelium. Antioxid Redox Signal 2019;30:1709–1730.

13. Kopacz A, Klóska D, Proniewski B, Cysewski D, Personnic N, Piechota-Polańczyk A, Kaczara P, Zakrzewska A, Forman HJ, Dulak J, Józkowicz A, Grochot-Przęczek A. Keap1 controls protein S-nitrosation and apoptosis-senescence switch in endothelial cells. Redox Biol 2020;28:101304.

14. Badi I, Burba I, Ruggeri C, Zeni F, Bertolotti M, Scopece A, Pompilio G, Raucci A. MicroRNA-34a Induces Vascular Smooth Muscle Cells Senescence by SIRT1 Downregulation and Promotes the Expression of Age-Associated Pro-inflammatory Secretory Factors. J Gerontol A Biol Sci Med Sci 2015;70:1304–1311.

15. Disayabutr S, Kim EK, Cha S-I, Green G, Naikawadi RP, Jones KD, Golden JA, Schroeder A, Matthay MA, Kukreja J, Erle DJ, Collard HR, Wolters PJ. miR-34 miRNAs Regulate Cellular Senescence in Type II Alveolar Epithelial Cells of Patients with Idiopathic Pulmonary Fibrosis – PubMed https://pubmed.ncbi.nlm.nih.gov/27362652/ (13 July 2023)

16. Ito T, Yagi S, Yamakuchi M. MicroRNA-34a regulation of endothelial senescence. Biochem Biophys Res Commun 2010;398:735–740.

17. Xu X, Chen W, Miao R, Zhou Y, Wang Z, Zhang L, Wan Y, Dong Y, Qu K, Liu C. miR-34a induces cellular senescence via modulation of telomerase activity in human hepatocellular carcinoma by targeting FoxM1/c-Myc pathway. Oncotarget 2015;6:3988–4004.

18. Boon RA, Iekushi K, Lechner S, Seeger T, Fischer A, Heydt S, Kaluza D, Tréguer K, Carmona G, Bonauer A, Horrevoets AJG, Didier N, Girmatsion Z, Biliczki P, Ehrlich JR, Katus HA, Müller OJ, Potente M, Zeiher AM, Hermeking H, Dimmeler S. MicroRNA-34a regulates cardiac ageing and function. Nature 2013;495:107–110.

19. Choi YJ, Lin C-P, Ho JJ, He X, Okada N, Bu P, Zhong Y, Kim SY, Bennett MJ, Chen C, Ozturk A, Hicks GG, Hannon GJ, He L. miR-34 miRNAs provide a barrier for somatic cell reprogramming. Nat Cell Biol 2011;13:1353–1360.

20. Zhong L, He X, Song H, Sun Y, Chen G, Si X, Sun J, Chen X, Liao W, Liao Y, Bin J. METTL3 Induces AAA Development and Progression by Modulating N6-Methyladenosine-Dependent Primary miR34a Processing. Mol Ther Nucleic Acids 2020;21:394–411.

21. Zalewski DP, Ruszel KP, Stępniewski A, Gałkowski D, Bogucki J, Komsta Ł, Kołodziej P, Chmiel P, Zubilewicz T, Feldo M, Kocki J, Bogucka-Kocka A. Dysregulation of microRNA Modulatory Network in Abdominal Aortic Aneurysm. J Clin Med 2020;9:1974.

22. Itoh K, Chiba T, Takahashi S, Ishii T, Igarashi K, Katoh Y, Oyake T, Hayashi N, Satoh K, Hatayama I, Yamamoto M, Nabeshima Y. An Nrf2/small Maf heterodimer mediates the induction of phase II detoxifying enzyme genes through antioxidant response elements. Biochem Biophys Res Commun 1997;236:313–322.

23. Zahreddine R, Davezac M, Smirnova N, Buscato M, Lhuillier E, Lupieri A, Solinhac R, Vinel A, Vessieres E, Henrion D, Renault M-A, Gadeau A-P, Flouriot G, Lenfant F, Laffargue M, Métivier R, Arnal J-F, Fontaine C. Tamoxifen Accelerates Endothelial Healing by Targeting ERα in Smooth Muscle Cells. Circ Res 2020;127:1473–1487.

24. Bar A, Kieronska-Rudek A, Proniewski B, Suraj-Prażmowska J, Czamara K, Marczyk B, Matyjaszczyk-Gwarda K, Jasztal A, Kuś E, Majka Z, Kaczor A, Kurpińska A, Walczak M, Pieterman EJ, Princen HMG, Chlopicki S. In Vivo Magnetic Resonance Imaging-Based Detection of Heterogeneous Endothelial Response in Thoracic and Abdominal Aorta to Short-Term High-Fat Diet Ascribed to Differences in Perivascular Adipose Tissue in Mice. J Am Heart Assoc 2020;9:e016929.

25. Targosz-Korecka M, Jaglarz M, Malek-Zietek KE, Gregorius A, Zakrzewska A, Sitek B, Rajfur Z, Chlopicki S, Szymonski M. AFM-based detection of glycocalyx degradation and endothelial stiffening in the db/db mouse model of diabetes. Sci Rep 2017;7:15951.

26. Menghini R, Casagrande V, Cardellini M, Martelli E, Terrinoni A, Amati F, Vasa-Nicotera M, Ippoliti A, Novelli G, Melino G, Lauro R, Federici M. MicroRNA 217 modulates endothelial cell senescence via silent information regulator 1. Circulation 2009;120:1524–1532.

27. Alexander Y, Osto E, Schmidt-Trucksäss A, Shechter M, Trifunovic D, Duncker DJ, Aboyans V, Bäck M, Badimon L, Cosentino F, De Carlo M, Dorobantu M, Harrison DG, Guzik TJ, Hoefer I, Morris PD, Norata GD, Suades R, Taddei S, Vilahur G, Waltenberger J, Weber C, Wilkinson F, Bochaton-Piallat M-L, Evans PC. Endothelial function in cardiovascular medicine: a consensus paper of the European Society of Cardiology Working Groups on Atherosclerosis and Vascular Biology, Aorta and Peripheral Vascular Diseases, Coronary Pathophysiology and Microcirculation, and Thrombosis. Cardiovasc Res 2021;117:29–42.

28. Kuhlencordt PJ, Gyurko R, Han F, Scherrer-Crosbie M, Aretz TH, Hajjar R, Picard MH, Huang PL. Accelerated atherosclerosis, aortic aneurysm formation, and ischemic heart disease in apolipoprotein E/endothelial nitric oxide synthase double-knockout mice. Circulation 2001;104:448–454.

29. Medina F, Haro J de, Florez A, Acin F. Relationship between endothelial dependent vasodilation and size of abdominal aortic aneurysms. Ann Vasc Surg 2010;24:752–757.

30. Lee R, Bellamkonda K, Jones A, Killough N, Woodgate F, Williams M, Cassimjee I, Handa A, Oxford Abdominal Aortic Aneurysm Study. Flow Mediated Dilatation and Progression of Abdominal Aortic Aneurysms. Eur J Vasc Endovasc Surg 2017;53:820–829.

31. Si Y, Chu H, Zhu W, Xiao T, Shen X, Fu Y, Xu R, Jiang H. Concentration-dependent effects of rapamycin on proliferation, migration and apoptosis of endothelial cells in human venous malformation. Exp Ther Med 2018;16:4595–4601.

32. Raucci A, Macrì F, Castiglione S, Badi I, Vinci MC, Zuccolo E. MicroRNA-34a: the bad guy in age-related vascular diseases. Cell Mol Life Sci 2021;78:7355–7378.

33. Kaller M, Liffers S-T, Oeljeklaus S, Kuhlmann K, Röh S, Hoffmann R, Warscheid B, Hermeking H. Genome-wide characterization of miR-34a induced changes in protein and mRNA expression by a combined pulsed SILAC and microarray analysis. Mol Cell Proteomics 2011;10:M111.010462.

34. Luo J, Su F, Chen D, Shiloh A, Gu W. Deacetylation of p53 modulates its effect on cell growth and apoptosis. Nature 2000;408:377–381.

35. Lu H, Du W, Ren L, Hamblin MH, Becker RC, Chen YE, Fan Y. Vascular Smooth Muscle Cells in Aortic Aneurysm: From Genetics to Mechanisms. Journal of the American Heart Association 2021;10:e023601.

36. DeRoo E, Stranz A, Yang H, Hsieh M, Se C, Zhou T. Endothelial Dysfunction in the Pathogenesis of Abdominal Aortic Aneurysm. Biomolecules 2022;12:509.

37. Segers VFM, Bringmans T, De Keulenaer GW. Endothelial dysfunction at the cellular level in three dimensions: severity, acuteness, and distribution. Am J Physiol Heart Circ Physiol 2023;325:H398–H413.

38. Quaschning T, Voss F, Relle K, Kalk P, Vignon-Zellweger N, Pfab T, Bauer C, Theilig F, Bachmann S, Kraemer-Guth A, Wanner C, Theuring F, Galle J, Hocher B. Lack of endothelial nitric oxide synthase promotes endothelin-induced hypertension: lessons from endothelin-1 transgenic/endothelial nitric oxide synthase knockout mice. J Am Soc Nephrol 2007;18:730–740.

39. Atochin DN, Huang PL. Endothelial nitric oxide synthase transgenic models of endothelial dysfunction. Pflugers Arch 2010;460:965–974.

40. Pimiento JM, Maloney SP, Tang PCY, Muto A, Westvik TS, Fitzgerald TN, Fancher TT, Tellides G, Dardik A. Endothelial nitric oxide synthase stimulates aneurysm growth in aged mice. J Vasc Res 2008;45:251–258.

41. Brady AR, Thompson SG, Fowkes FGR, Greenhalgh RM, Powell JT. Abdominal Aortic Aneurysm Expansion. Circulation 2004;110:16–21.

42. Kolluru GK, Bir SC, Kevil CG. Endothelial Dysfunction and Diabetes: Effects on Angiogenesis, Vascular Remodeling, and Wound Healing. Int J Vasc Med 2012;2012:918267.

43. Cafueri G, Parodi F, Pistorio A, Bertolotto M, Ventura F, Gambini C, Bianco P, Dallegri F, Pistoia V, Pezzolo A, Palombo D. Endothelial and smooth muscle cells from abdominal aortic aneurysm have increased oxidative stress and telomere attrition. PLoS One 2012;7:e35312.

44. Franck G, Dai J, Fifre A, Ngo S, Justine C, Michineau S, Allaire E, Gervais M. Reestablishment of the endothelial lining by endothelial cell therapy stabilizes experimental abdominal aortic aneurysms. Circulation 2013;127:1877–1887.

45. Yang X, Xu C, Yao F, Ding Q, Liu H, Luo C, Wang D, Huang J, Li Z, Shen Y, Yang W, Li Z, Yu F, Fu Y, Wang L, Ma Q, Zhu J, Xu F, Cong X, Kong W. Targeting endothelial tight junctions to predict and protect thoracic aortic aneurysm and dissection. Eur Heart J 2023;44:1248–1261.

46. Hoshina K, Sho E, Sho M, Nakahashi TK, Dalman RL. Wall shear stress and strain modulate experimental aneurysm cellularity. Journal of Vascular Surgery 2003;37:1067.

47. Kiosses WB, McKee NH, Kalnins VI. Evidence for the migration of rat aortic endothelial cells toward the heart. Arterioscler Thromb Vasc Biol 1997;17:2891–2896.

48. Roostalu U, Aldeiri B, Albertini A, Humphreys N, Simonsen-Jackson M, Wong JKF, Cossu G. Distinct Cellular Mechanisms Underlie Smooth Muscle Turnover in Vascular Development and Repair. Circ Res 2018;122:267–281.

49. Foteinos G, Hu Y, Xiao Q, Metzler B, Xu Q. Rapid endothelial turnover in atherosclerosis-prone areas coincides with stem cell repair in apolipoprotein E-deficient mice. Circulation 2008;117:1856–1863.

50. Li Y, Liu Z, Han X, Liang F, Zhang Q, Huang X, Shi X, Huo H, Han M, Liu X, Zhu H, He L, Shen L, Hu X, Wang J, Wang Q-D, Smart N, Zhou B, He B. Dynamics of Endothelial Cell Generation and Turnover in Arteries During Homeostasis and Diseases. Circulation 2024;149:135–154.

51. Lukowski SW, Patel J, Andersen SB, Sim S-L, Wong HY, Tay J, Winkler I, Powell JE, Khosrotehrani K. Single-Cell Transcriptional Profiling of Aortic Endothelium Identifies a Hierarchy from Endovascular Progenitors to Differentiated Cells. Cell Rep 2019;27:2748–2758.e3.

52. McDonald AI, Shirali AS, Aragón R, Ma F, Hernandez G, Vaughn DA, Mack JJ, Lim TY, Sunshine H, Zhao P, Kalinichenko V, Hai T, Pelegrini M, Ardehali R, Iruela-Arispe ML. Endothelial Regeneration of Large Vessels Is a Biphasic Process Driven by Local Cells with Distinct Proliferative Capacities. Cell Stem Cell 2018;23:210–225.e6.

53. Shirali AS, Romay MC, McDonald AI, Su T, Steel ME, Iruela-Arispe ML. A multi-step transcriptional cascade underlies vascular regeneration in vivo. Sci Rep 2018;8:5430.

54. Birdsey GM, Shah AV, Dufton N, Reynolds LE, Osuna Almagro L, Yang Y, Aspalter IM, Khan ST, Mason JC, Dejana E, Göttgens B, Hodivala-Dilke K, Gerhardt H, Adams RH, Randi AM. The Endothelial Transcription Factor ERG Promotes Vascular Stability and Growth through Wnt/β-Catenin Signaling. Dev Cell 2015;32:82–96.

55. Dryden NH, Sperone A, Martin-Almedina S, Hannah RL, Birdsey GM, Khan ST, Layhadi JA, Mason JC, Haskard DO, Göttgens B, Randi AM. The Transcription Factor Erg Controls Endothelial Cell Quiescence by Repressing Activity of Nuclear Factor (NF)-κB p65. J Biol Chem 2012;287:12331–12342.

56. Zhan J, Qin S, Lu L, Hu X, Zhou J, Sun Y, Yang J, Liu Y, Wang Z, Tan N, Chen J, Zhang C. miR-34a is a common link in both HIV- and antiretroviral therapy-induced vascular aging. Aging (Albany NY) 2016;8:3298–3310.

57. Sangiorgi’ ‘Giuseppe, Martelli’ ‘Eugenio, Tolva’ ‘Valerio, Cotroneo’ ‘Attilio, Micari’ ‘Antonio, Luca’ ‘Fabio De, Cereda’ ‘Alberto, Trimarchi’ ‘Santi. Role of biochemical markers in the diagnosis and treatment of an aneurysm of the abdominal aorta https://www.escardio.org/Journals/E-Journal-of-Cardiology-Practice/role-of-biochemical-markers-in-the-diagnosis-and-treatment-of-an-aneurysm-of-the (12 December 2023)

58. Klopf J, Brostjan C, Neumayer C, Eilenberg W. Neutrophils as Regulators and Biomarkers of Cardiovascular Inflammation in the Context of Abdominal Aortic Aneurysms. Biomedicines 2021;9:1236.

59. Schmidt-Ott KM, Kagiyama S, Phillips MI. The multiple actions of angiotensin II in atherosclerosis. Regulatory Peptides 2000;93:65–77.

60. Vizcaíno JA, Csordas A, Toro N del-, Dianes JA, Griss J, Lavidas I, Mayer G, Perez-Riverol Y, Reisinger F, Ternent T, Xu Q-W, Wang R, Hermjakob H. 2016 update of the PRIDE database and its related tools. Nucleic Acids Res 2016;44:D447–456.

